# Angular head velocity cells within brainstem nuclei projecting to the head direction circuit

**DOI:** 10.1101/2023.03.29.534808

**Authors:** Jalina A. Graham, Julie R. Dumont, Shawn S. Winter, Joel E. Brown, Patrick A. LaChance, Carly C. Amon, Kara B. Farnes, Ashlyn J. Morris, Nicholas A. Streltzov, Jeffrey S. Taube

## Abstract

An animal’s perceived sense of orientation depends upon the head direction (HD) system found in several limbic structures and depends upon an intact peripheral vestibular labyrinth. However, how the vestibular system influences the generation, maintenance, and updating of the HD signal remains poorly understood. Anatomical and lesion studies point towards three key brainstem nuclei as being potential critical components in generating the HD signal: nucleus prepositus hypoglossi (NPH), supragenual nucleus (SGN), and dorsal paragigantocellularis reticular nuclei (PGRNd). Collectively, these nuclei are situated between the vestibular nuclei and the dorsal tegmental and lateral mammillary nuclei, which are thought to serve as the origin of the HD signal. To test this hypothesis, extracellular recordings were made in these areas while rats either freely foraged in a cylindrical environment or were restrained and rotated passively. During foraging, a large subset of cells in all three nuclei exhibited activity that correlated with changes in the rat’s angular head velocity (AHV). Two fundamental types of AHV cells were observed: 1) symmetrical AHV cells increased or decreased their neural firing with increases in AHV regardless of the direction of rotation; 2) asymmetrical AHV cells responded differentially to clockwise (CW) and counter-clockwise (CCW) head rotations. When rats were passively rotated, some AHV cells remained sensitive to AHV whereas others had attenuated firing. In addition, a large number of AHV cells were modulated by linear head velocity. These results indicate the types of information conveyed in the ascending vestibular pathways that are responsible for generating the HD signal.

**Significance Statement:** Extracellular recording of brainstem nuclei (nucleus prepositus hypoglossi, supragenual nucleus, and dorsal paragigantocellularis reticular nucleus) that project to the head direction circuit identified different types of angular head velocity (AHV) cells while rats freely foraged in a cylindrical environment. The firing of many cells was also modulated by linear velocity. When rats were restrained and passively rotated some cells remained sensitive to AHV, whereas others had attenuated firing. These brainstem nuclei provide critical information about the rotational movement of the rat’s head in the azimuthal plane.

## Introduction

The ability to navigate successfully is critical for survival and relies on neural circuitry that provides information about spatial location and orientation. For instance, an animal’s perceived sense of direction depends upon the head direction (HD) system, which is comprised of neurons that fire when an animal’s head points a particular direction in the environment. Each HD cell is tuned to a specific angle, referred to as the cell’s preferred firing direction (PFD), and the population of HD cells encode for all directions (Taube, 2007). HD cells are found in brain areas primarily along Papez’ circuit and the HD signal is thought to be generated across the reciprocal connections between the dorsal tegmental nucleus (DTN) and lateral mammillary nucleus (LMN) (Sharp et al., 2001a; Bassett et al., 2007).

In the absence of spatial landmarks directional heading can be determined by integrating changes in angular head velocity (AHV) when the initial directional heading is known. Changes in AHV can be derived from vestibular, proprioceptive, or motor efference information. Most computational models have argued for an important role for the vestibular labyrinth, which tracks both angular and linear velocity of the head (McNaughton et al., 1991; Redish et al. 1996). Indeed, damage to the peripheral vestibular system disrupts direction-specific firing in HD cells (Stackman and Taube, 1997; Muir et al., 2009; Valerio and Taube, 2016). Activity in DTN is also primarily related to AHV, along with a small percentage of HD modulated cells (Bassett and Taube, 2001; Sharp et al., 2001b). When compared to vestibular afferents, which respond to one turn direction (CW or CCW), many cells in the DTN show bilaterally symmetric responses to head turns with firing rates increasing linearly in both turn directions. These symmetric cells have been predicted to play an important role in the ring attractor network (Stratton e al., 2010), but it is currently unknown how the vestibular response profile in DTN is created.

While at least one study has found a sparse connection between the vestibular nuclei and DTN (Liu et al., 1984), vestibular activity in DTN is likely driven by direct input from a complex network of intermediate brainstem nuclei that receive information from the medial vestibular nucleus (MVN). These nuclei include the nucleus prepositus hypoglossi (NPH), dorsal paragigantocellularis reticular nucleus (PGRNd), and supragenual nucleus (SGN). Notably, the MVN receives information from the horizontal semicircular canals and is sensitive to head turns in yaw (Uchino et al., 2005). Figure 1 shows the extensive connectivity between these nuclei and how they ultimately project to the DTN and LMN (Liu et al., 1984; Biazoli et al., 2006). Lesions or inactivation of either SGN or NPH eliminated direction-specific firing in anterodorsal thalamic (ADN) neurons and also impaired homing behavior (Clark et al., 2012; Butler et al., 2017). In contrast to the NPH and SGN, the contribution of the PGRNd to the HD signal remains unknown.

**Figure 1.**
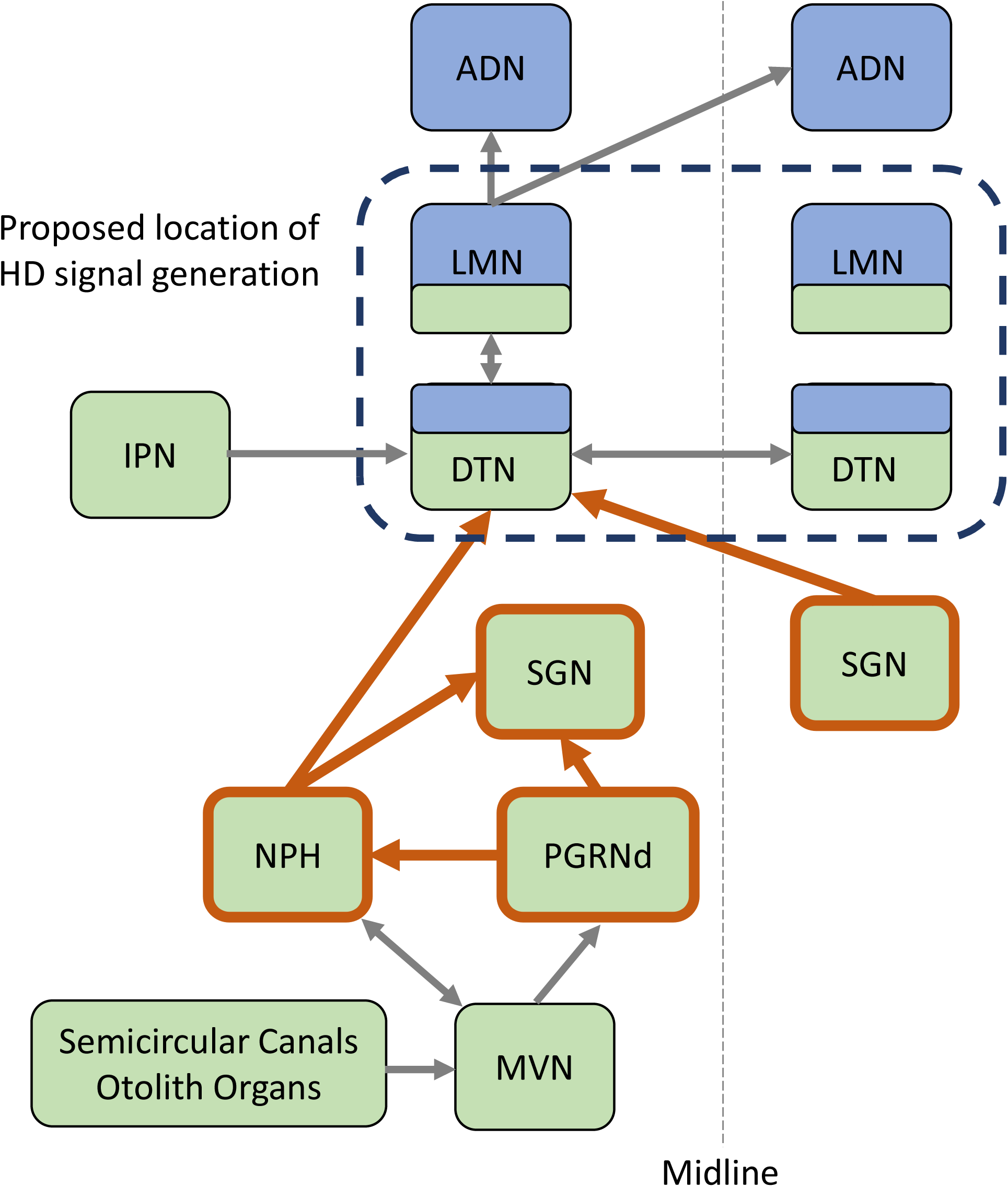
Circuitry showing the major connections of the vestibular inputs into the HD network (highlighted by orange borders). Green-colored squares represent areas where AHV cells have been identified and blue-colored squares indicate areas where HD cells have been identified. Brain areas discussed in this study are all afferent to the DTN and their interconnections in the brainstem are highlighted in orange. Abbreviations: ADN: anterior dorsal nucleus of the thalamus; DTN: dorsal tegmental nucleus of Gudden; IPN: interpeduncular nucleus; LMN: lateral mammillary nucleus; MVN: medial vestibular nucleus; NPH: nucleus prepositus hypoglossi; PGRNd: paragigantocellularis reticular nucleus dorsal; SGN: supragenual nucleus.

To further understand how the NPH, SGN, and PGRNd may contribute to the HD system, single-unit recordings were made in these areas while rats freely foraged or were restrained and passively rotated. The rationale for examining rats during passive rotation comes from evidence that vestibular neurons in monkeys increase their firing when the animal is passively rotated, but display marked attenuation during voluntarily movements (McCrea et al., 1999; Roy and Cullen, 2001, 2004). These findings would suggest that during voluntary movement (free-foraging), the AHV signal would be attenuated and incapable of updating the animal’s perceived directional heading. This rationale, however, contradicts the studies showing that the HD signal depends upon vestibular inputs. Because of this conundrum (see Shinder and Taube, 2014; Cullen and Taube, 2017), a subset of AHV cells was examined under passive head-fixed restraint conditions. We report that similar to recordings in the DTN, all three brainstem nuclei contain cells that exhibit firing correlated with AHV and likely provide critical information for updating the HD signal following a head turn.

## Methods

### Subjects

The subjects were 39 female Long-Evans rats (Envigo), weighing 225-350 g at the time of surgery. The rats were housed individually following surgery, and were on a 12:12h light-dark cycle. To increase spatial sampling rats performed a continuous foraging for pellets task. Rats were initially food restricted to no less than 85% of their free-feeding weight in order to motivate them to perform the task. All rats had *ad libitum* access to water. All procedures were performed in compliance with institutional standards set forth by the *National Institutes of Health Guide for the Care and Use of Laboratory Animals* and the Society for Neuroscience.

### Surgery and electrodes

Rats were anaesthetized with either isoflurane gas or a ketamine-xylazine mixture (2ml/kg intramuscular) and placed into a stereotaxic frame (David Kopf instruments). A craniotomy of ~0.5 mm diameter was drilled over the brainstem targets. The PGRNd lies just ventral to the NPH; thus, the coordinates for PGRNd were the same as for NPH and the electrode array was advanced further ventrally when recording from the NPH. The coordinates with respect to lambda using the Paxinos and Watson (2005) rat brain atlas were: anterior NPH: anterior-posterior (A-P): 3.5-3.7 mm, medial-lateral (M-L): ± 0.3-0.5 mm, dorsal-ventral from the cortical surface (D-V): 6.0-7.0 mm; posterior NPH: A-P: 4.9 mm, M-L: ± 0.45 mm, D-V: 6.0 mm; SGN: A-P: 3.5 mm, M-L: ± 0.45-0.50 mm, D-V: 6.0-6.5 mm. There were no differences found between cells recorded in the anterior versus posterior NPH and cells from both coordinates were therefore grouped together in the analyses. For all coordinates, the electrodes were implanted into the brainstem at −10° angle along the anterior-posterior axis (i.e., the electrode tips were pointed rostrally towards the rat’s nose along the anterior-posterior axis) to avoid the transverse sinus.

Electrodes consisted of either a bundle of 4 or 10 single wires (n=14), an array of 3 or 8 stereotrodes (n=15), or 1 tetrode (n=10). The single wire electrodes were built as previously described (Kubie, 1984; Taube, 1995) and consisted of either a bundle of four or ten 25 μm nichrome wires threaded through a 26-gauge stainless steel cannula and connected to a modified Augat plug. Stereotrodes and tetrodes were made by spinning 17 μm nichrome wires into a compact bundle composed of either 2 (stereotrodes) or 4 wires (tetrodes)(Winter et al., 2015). These wires were threaded through a smaller 32-gauge cannula that was connected to a modified Mill-Max plug. In all cases, electrode design was chosen to minimize tissue distortion/damage while maximizing cell yield in these small-sized brain areas. Five to 8 stainless-steel screws were placed along the top of the skull to help secure the electrode arrays to the skull using dental acrylic. In addition, some animals (n=9) had a post (4.76 mm square, ~18 mm long, item-8737K12, Teflon® PTFE, McMaster-Carr) affixed to their skull with dental acrylic that could be used to head-fix the rat’s head to a restraint device (see Shinder and Taube, 2011). This set-up was used to prevent the rats from moving their heads during a passive movement condition (described below). The rats that had a head-post implanted had 6 additional screws placed along the temporal and the most lateral parts of the occipital bone plates. Once the electrode arrays were lowered into the brain, they were secured to the skull with dental acrylic, but could be manipulated in the dorsal/ventral plane by turning three drive screws that were part of the electrode assembly. Rats were given the analgesic ketoprofen (3-5 mg/kg, intramuscular) post-operatively for two consecutive days and were given one week of recovery before commencing recording.

### Apparatuses and Training

Rats were initially placed on a food-restricted diet and trained to retrieve randomly scattered sucrose food pellets on the floor of either a cylindrical (n=36) or square (n=3) enclosure. Once the rat spent the majority of their time moving through the cylinder they underwent surgery for the implantation of electrodes. Two different apparatuses were used depending on whether the recordings were made in an active session (free-foraging) or a passive session.

### Active Sessions

During free-foraging, rats were placed inside a gray-painted cylindrical wooden enclosure (76 cm in diameter, 51 cm high) with disposable gray photographic backdrop paper covering the floor. Floor-to-ceiling curtains surrounded the cylinder, which eliminated the use of distal visual cues. A white sheet of cardboard that was 51 cm high and occupying 100° of arc was attached to the inside wall of the cylinder. This white cue card served as the only intentional visual landmark cue for the animal when it was inside the cylinder. Eight DC lights, suspended overhead and arranged uniformly in a circle (diameter 71 cm), provided the illumination. A color video camera (SONY XC-711) was centered above the cylinder and 3 m above the floor. A ceiling-mounted food dispenser automatically ejected sucrose pellets (20 mg, Dustless Precision Pellets, Bio-Serv) on a 30 s random interval schedule. During the free foraging sessions rats searched for and consumed the sucrose pellets. These sessions were usually 16 min long although a small number of sessions were 8 min long. The sessions conducted in the square enclosure (1 m × 1 m, 50 cm high) used the same procedures as above; a white cue card (61 cm × 50 cm) was centered on one wall of the square.

### Passive sessions

For the passive condition, we conducted one of two types of sessions because different types of passive movements have been applied in the past (Taube, 1995; Sharp et al., 2001b; Shinder and Taube, 2011). For 9 sessions, the rat was head-fixed restrained and placed on a rotating platform (48 cm long; 9 cm wide; 22 cm high), forming a lazy Susan as previously described by Shinder and Taube (2011; see Fig. 1). The platform was positioned inside the cylinder. Rats were gradually habituated to the passive restraint apparatus. In the first stage of habituation (Stage 1), the rats were hand-held in a towel for a minimum of three 10 min sessions. A session was considered successful if the rat remained still. For Stage 2, the rats were wrapped in the towel, which was held in place with zip ties, and placed inside a plastic tube (6 cm in diameter) on a lazy Susan for another three sessions (minimum). The plastic tube did not extend rostrally beyond the towel, and therefore was not in the way of the rat’s head. Once acclimated to the tube, rats received their electrode implantation. Following 5-7 d post-operative recovery, the rats were reintroduced to Stage 2 for a further three sessions (or more if needed). Finally, in the third and final stage of habituation, the rat’s head was restrained by attaching the implanted post on the rat’s head to a mounted bar that was attached to the passive apparatus via a coupling sleeve. The axis of rotation was always aligned with the animal’s intra-aural axis of the head (i.e., rotation about the head’s dorsal-ventral axis). The tube was held in place on the platform of the lazy Susan with the use of bungee cords so that the rat’s head remained aligned along the longitudinal axis of the restraint apparatus. This procedure reduced pressure on the animal’s head, neck, or body. Because the entire passive apparatus fit within the cylindrical arena, the same visual cues were available to the rat during active and passive sessions. The experimenter, who was stationed outside the cylinder, would reach in and manually rotate the platform.

Head-fixed passive sessions were ~8 min in duration and consisted of the following sequence: 1) continuous CCW motion, 2) sinusoidal (back-and-forth) motion, 3) continuous CW motion, 4) sinusoidal motion, 5) 90° CCW steps (abrupt stop every 90° rotation), 6) 90° CW steps, 7) 45° CCW steps, and 8) 45° CW steps. Rotational speeds contained a large range of movement velocities, but generally varied between 0-270°/s. During these sweeps the rat was gradually turned through 360° in order to sample all directional orientations. Following the passive session, a second 16 min active session was recorded to verify the cell’s waveform isolation and whether any changes that occurred under passive restraint may have contributed to changes in the cell’s firing characteristics.

For the other type of passive session (n=18), the rat was wrapped in a towel such that only its head and associated headstage were exposed outside the towel. Then the towel-wrapped rat was held in the experimenter’s hands and moved back and forth in 90-150° sweeps just above the floor of the cylinder. The speed of the rotations varied between 0-300°/s. Recording sessions lasted 1 min.

### Isolation and recording of single-unit activity

Activity on each electrode wire, stereotrode, or tetrode was examined across daily screening sessions while the rats foraged for food pellets within the cylindrical enclosure. A cell was recorded when single-unit waveform activity was isolated above background activity. Electrical signals from the electrode wires were passed through a field-effect transistor in a source-follower configuration. For recordings made with single wires, the signals were then amplified by a factor of 20K (Grass Instruments), band-pass filtered (300-10,000 Hz) and passed through a dual-window discriminator (BAK Electronics) for spike discrimination. The occurrence of spikes was then timestamped from the start of the session and the timestamps were sent to a computer for off-line analysis with the video data. The video data involved tracking the position and directional orientation of the rat using red and green light-emitting diodes (LEDs), which were secured to the animal’s headstage ~9 cm apart. The red LED was positioned above the rat’s head and the green LED was positioned above the rat’s back. The X and Y coordinates of each LED position was recorded at a sampling rate of 60 Hz using an automated video tracking system (Eberle Electronics), and the rat’s HD and location were determined by using the relative positions of the two LEDs.

Signals originating from the stereotrodes or tetrodes were pre-amplified by unity-gain operational amplifiers on an HS-27 headstage, then further amplified (Neuralynx-8 amplifier) and then band-pass filtered (600-6,000 Hz) using an ERP-27 system (Neuralynx). When the neural signals crossed a pre-set amplitude threshold (~30-60 μV), they were timestamped and digitized at 32 KHz for 1 msec. The waveform characteristics were then analyzed offline using SpikeSort3D (Neuralynx). The rat’s location and directional heading were also tracked in a similar manner as above using two differently colored LEDs.

Each rat was hand carried into the recording room, and no attempt was made to disorient the rat when it was placed in the cylinder, except for the one series of sessions when the cue card was rotated (see Fig. 9B below). For all electrode types, the wires were advanced at the end of each session (30-115 µm). Screening for cells took place approximately daily over the course of 2-4 months after which the rats were euthanized and perfused with formalin.

## Data Analyses

### Angular head velocity

AHVs were determined using methods described previously (Taube, 1995; Stackman and Taube, 1998; Bassett and Taube, 2001). Briefly, the rat’s HD was first determined from the relative position of the two LEDs, and sorted into sixty 6° bins. The average firing rate for each bin (i.e., summed number of spikes in a given bin divided by time spent in the bin) was calculated for the duration of the recording session. The AHVs were then computed from the HD values creating a HD by time function that was smoothed across five time points using the following function: n = (n_t-2_ + n_t-1_ + n + n_t+1_ + n_t+2_)/5. From the smoothed HD × time function, we took episodes of five samples and calculated the best-fit slope (derivative) and this value was defined as the AHV for the center value of the five samples. This procedure was repeated for all samples and then the AHV from all samples, along with the number of corresponding spikes for that sample, were summed to create a firing rate vs. AHV plot based on 6°/s bins.

Because high AHVs must necessarily pass through the range of lower AHVs, there is an inherent sampling bias towards the lower AHVs, resulting in increased variability accompanied with fewer samples at the higher velocities. To minimize the effect of this sampling bias, AHV bins that contained fewer than 30 samples (0.5 s of recording time) and all bins above 204°/sec were excluded. The mean velocity and mean firing rate for each 6°/sec AHV bin were calculated and plotted into a firing rate by AHV scattergram for each session. From the firing rate-AHV functions, several parameters were calculated: 1) baseline firing rate defined as the mean firing rate when the rat’s AHV was near zero (between ± 6°/sec bins), 2) the Pearson correlation coefficient for the linear best-fit lines of the CW and CCW functions; CCW turns were defined as positive values; CW turns as negative values, and 3) the best-fit lines’ slopes.

## Cell classification

There are a number of possible methods for classifying cells as sensitive to AHV including 1) meeting a certain correlation and slope threshold on a firing rate vs. AHV plot, 2) passing a shuffled test for the same criteria, 3) performing a generalized linear model (GLM) on the data. Here, we have performed all three methods, but have selected the first method above as we found this approach to best approximate what we observed with respect to the cells’ firing rate vs. AHV tuning curves (see discussion of this issue in Results).

The scattergram for each firing rate vs. AHV plot was divided into four AHV ranges (CW: 0-90°/s, CW: 90-204°/s, CCW: 0-90°/s, CCW: 90-204°/s). For each range of points, we calculated the best-fit line for those points and its corresponding Pearson’s correlation (*r*) value and slope. Because most of the rat’s angular turns were ≤ 90°/s, this portion of the AHV tuning curve had the best sampling and was the most reliable. We therefore used correlation and slope values in the analyses below from this range of AHVs rather than the entire 0-204°/s range. We note, however, that 1) any cell that had a significant correlation and slope in the upper range (90-204°/s), also exhibited a significant correlation in the lower range values (0-90°/s), and 2) the results did not change significantly if a different range of AHV values were used for the analyses (e.g., 0-150°/s).

Cells were classified as AHV if they met the following two criteria: 1) one of the two 0-90°/s ranges for CW or CCW values contained a correlation ≥ 0.5 and 2) the absolute value of the slope for the best-fit line was ≥ 0.025 spikes/°. We also tested each cell that passed these thresholds against a shuffled test and with a generalized linear model (see below). For the shuffle test, the spike data was shifted randomly by at least 5 s relative to the position tracking data 500 times. Then, for each shuffle an AHV vs. firing rate curve was generated and best-fit line values calculated for the two AHV ranges. To pass the shuffle test the unshuffled based plot had to rank above the 95^th^ percentile of the 500 values.

AHV cells were divided into AHV cell types based on a normalized turn bias score defined as:

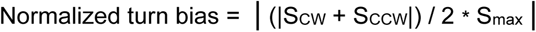

where S_CW_ and S_CCW_ are the CW and CCW slopes between 0 to ± 90°/s, respectively; S_max_ is the larger of the two slopes’ absolute values. Using this measure, cells that are perfectly symmetrical between their CW and CCW tuning curves will have normalized turn bias values near zero while perfectly asymmetric cells will have values near 1. A cell was classified as symmetric when its normalized turn bias was ≤ 0.3, asymmetric when its normalized turn bias was ≥ 0.7, and asymmetric-unresponsive for values in between.

Of the non-AHV cells, 28% were classified as both linear velocity and AHV by the GLM, but failed to pass shuffle or slope/correlation cutoffs, suggesting that these cells may be AHV and linear velocity modulated, but failed to pass threshold. However, the pattern of statistical results remains the same if thresholding criteria or GLM are used. Some non-AHV cells (2.4%) could be classified as linear velocity alone, but all other non-AHV cells had no clear responses related to self-motion.

## Linear velocity

For linear velocity we first computed the instantaneous speed of the animal for each 1/60^th^ sec sample, by fitting a best-fit line over a 5-sample window for the x and y dimensions (Bassett et al., 2007). The slopes of the best-fit lines were defined as the change in the x and y dimensions, respectively, and the instantaneous speed for the center time point of the window was defined as the square root of x^2^ + y^2^. We then constructed a linear velocity tuning curve by sorting all samples based on 1 cm/s bins. The number of spikes for each sample were also sorted based on the sample’s LV. Next, the number of spikes in each linear velocity bin were summed and divided by the total time in that bin to yield an average firing rate for that linear velocity bin. A firing rate vs. linear velocity plot was then created and a best-fit linear line was computed between 0-30 cm/s. From this linear fit we calculated a Pearson’s *r* correlation coefficient and its corresponding slope. We classified a cell as modulated by linear velocity if the absolute value of its correlation ≥ 0.7, the absolute value of its slope ≥ 0.1, and the correlation and slope values passed the 95^th^ percentile of tuning curves generated from a shuffled time series (similar to the approach for AHV analyses).

### Cell classifications with a generalized linear model

Cells were classified as encoding up to four behavioral variables using 10-fold cross-validation with a Poisson generalized linear model (GLM; Hardcastle et al., 2017). The behavioral variables were: allocentric HD, 2D location, angular head velocity, and linear velocity. Briefly, for a given model, the firing rate vector *r* for a single cell over all time points was modeled as follows:

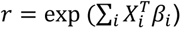

where *X* is a matrix containing animal state vectors for a single behavioral variable across time points *T*, β represents the parameter vector for that behavioral variable (similar to a tuning curve), and *i* indexes across behavioral variables included in the model. The parameter vectors for a given model are learned by maximizing the log-likelihood *l* of the real spike train *n* given the model’s estimated rate vector *r*:

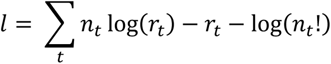

Where *t* indexes over time points. In order to avoid overfitting for the cross-validation procedure, an additional smoothing penalty *P* was added to the objective function which penalizes differences between adjacent bins of each parameter vector (similar to fused ridge regularization):

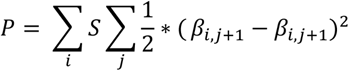

Here, *S* is a smoothing hyperparameter (20 for HD, AHV, and linear velocity; 2 for 2D location), *i* indexes over variables, and *j* indexes over response parameters for a given variable. Response parameters were estimated by minimizing (P – *l*) using SciPy’s *optimize.minimize* function. Thirty bins were used for allocentric head direction parameter vectors, ten bins were used for linear velocity (from 0 to 40 cm/s), 20 bins were used for AHV (from −200 to +200 deg/s), and 5 × 5 cm bins (20 × 20 for the 100 cm square, 14 × 14 for the 70 cm diameter cylinder) were used for 2D location.

For cross-validation, data for a session was split into training (9/10 of the session) and test (1/10 of the session) data (*k* = 10 folds). Parameter vectors were estimated by minimizing the objective function on the training data using the full model with all four variables. Drawing parameter estimates from the full model helps to reduce correlation artifacts between variables (Burgess et al., 2005) and makes models with different variable combinations more comparable. Log-likelihoods for models with all possible variable combinations were computed. This procedure was repeated until all portions of the data had been used as test data (10 folds).

To select the best model, the log-likelihood values from the best two-variable model were compared to those from the best one-variable model. If the two-variable model showed significant improvement from the one-variable model (using a one-sided Wilcoxon signed-rank test), then the best three-variable model was compared to the two-variable model, and so on. If the more complex model was not significantly better, the simpler model was chosen. If the chosen model performed significantly better than an intercept-only model, the chosen model was used as the cell’s classification. Otherwise, the cell was marked ‘unclassified’ (Hardcastle et al., 2017).

To assess the contribution of each variable to the firing of a cell, we optimized the cell’s chosen model based on data from the full recording session, and then computed the difference in several goodness-of-fit measures when removing a particular variable from the model. For example, if a cell were tuned to HD and AHV, we would compute the goodness-of-fit of the HD and AHV model, and then calculate the contribution of AHV by calculating the loss of goodness-of-fit when removing AHV from the model. One goodness-of-fit measurement was log-likelihood per spike (measured in bits/spike), which was calculated by dividing the log-likelihood of the model by the total number of spikes in the session. Another was a pseudo-R^2^ measure based on the log-likelihood (Cameron and Windmeijer, 1996):

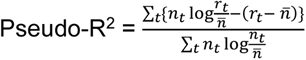

Where 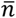 denotes the cell’s mean firing rate.

We also used two goodness-of-fit measures that required first binning the modeled firing rate vector and spike train into 300 ms bins. One of these was Pearson’s *r*, which assessed the correlation between the modeled rate vector and spike train. The other was explained variance (R^2^), computed as follows:

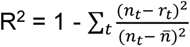

### Statistical procedures

Within region comparisons of the normalized turn bias scores for symmetrical and asymmetrical cells used independent sample t-tests. A two-way ANOVA was used for comparisons of numerical data, with Region (NPH, SGN, PGRNd) and Cell Type (Non-AHV, Symmetrical, Asymmetrical) or contralateral selectivity (Contralateral, Ipsilateral) as between subject factors. Tukey HSD post-hoc tests were used to explore significant main effects.

A Chi-square test was used to examine if the proportion of qualitative data (AHV cell type or contralateral selectivity) differed significantly across the three brainstem regions. Pairwise post-hoc Chi-square tests were conducted if the overall Chi-square test was significant to confirm the main differences between brain regions. Post-hoc Chi-square significance threshold was adjusted to 0.17 using a Bonferroni correction for multiple comparisons (0.05/3 comparisons).

### Histology

To facilitate localization of the recorded neurons, the electrode wires were only lowered ~225 μm past the depth at which the last AHV cell was recorded. Once the final depth of the electrode was reached, the rats were given an overdose of sodium pentabarbitol (100 mg/ml, intraperitoneal). The locations of two electrode wires were marked by passing weak anodal current (15 μA for 5s or 15s) in order to perform a Prussian blue reaction. The rats were perfused transcardially with 0.9% saline followed by 10% formalin. The brains were removed from the skull and post-fixed in 10% formalin for a minimum of 48 hr. Afterwards, 2% potassium ferrocyanide was added to the formalin solution for 24 hr. The brains were then transferred to 20-30% sucrose solution for at least 24 hr prior to being frozen and sectioned in the coronal plane at 30 μm slices on a cryostat. Sections throughout the extent of the targeted area (NPH/PGRNd or SGN) were mounted onto gelatin-coated slides. The slides were stained with thionin and examined under the microscope to determine the location of the marking lesion.

Lesions marked the final position of the wires. The position of the ventral-most point relative to the boundaries of the NPH, SGN, and PGRNd was used to estimate the portion of the screening record that could have fallen within these structures. Only the cells recorded within the dorsal-ventral extent of the NPH (~500 μm), SGN (~200 μm), and PGRNd (~800 μm) were used for analysis. When wires from the same electrode passed through different brainstem nuclei, only those that received the marking lesion were used to determine the location of the cells. In these cases, cells recorded from unmarked wires were excluded. Despite using this conservative approach, it remains possible some cells may have been recorded from other nuclei close to the border of our three regions of interest (e.g., central grey surrounding the SGN) due to the small size of these nuclei.

## Results

### Regional comparisons of AHV modulation

We recorded 523 brainstem neurons from 39 rats while the rats freely foraged for food pellets in a cylindrical or square-shaped arena. Figures 2 and 3 show examples of electrode tracks from recording wires that passed through the areas of interest as well as a summary of estimated locations for each electrode track and cell. Approximately equal numbers of cells were recorded in all areas: NPH (16 rats, 178 cells), SGN (20 rats, 158 cells), PGRNd (15 rats, 187 cells). Because the PGRNd is situated just ventral to the NPH, electrodes passed through both NPH and PGRNd in 12 rats. AHV × firing rate turning curves were constructed for each cell and from these plots we calculated the Pearson *r* values and slopes for each turn direction, as well as an estimate of the cell’s baseline firing rate when the rat is generally not turning its head (i.e., mean firing rate for the 0-6°/s bins in the CW and CCW directions). Correlation and slope values were obtained from AHV values in the range of 0-90°/s, since the majority of behavioral sampling occurred in this range, with rats spending on average 74% of the time moving their heads at ≤ 90°/s ( 1^st^ through 3^rd^ quartiles: 70 - 78%). Using the absolute values of the correlation and slope measures for CW and CCW turns, we defined the maximum correlation (max *r*) and the maximum slope (max slope) as the larger of the two values when comparing CW and CCW values.

**Figure 2.**
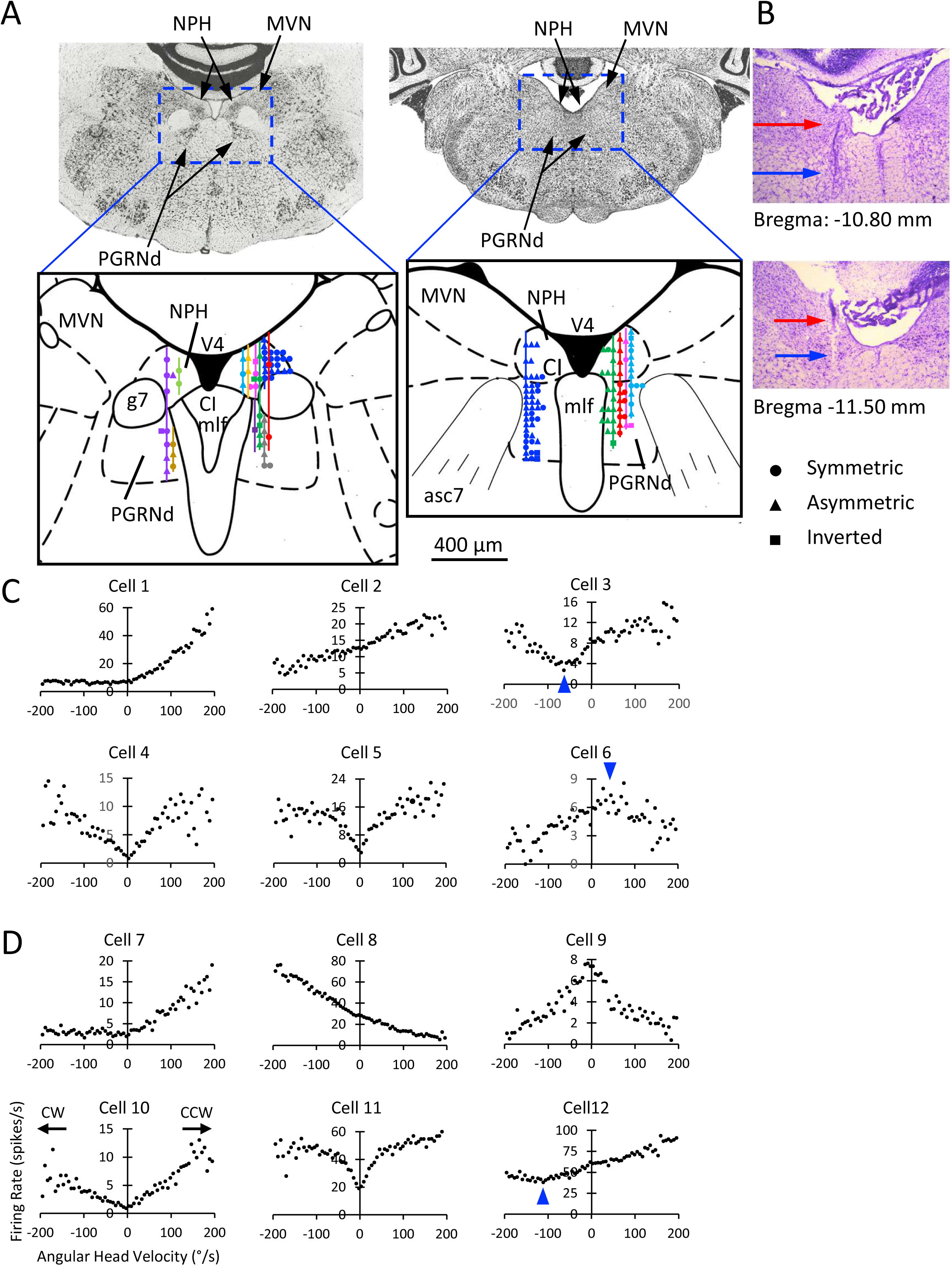
**A)** Photographs and schematic illustrations of coronal sections showing the location of NPH and PGRNd within the brainstem at −10.80 mm (left) and −11.50 mm (right) posterior to bregma. Each color track represents a different animal. **B)** Corresponding photomicrographs of example coronal sections stained with thionin at the level of anterior (top) and posterior NPH/PGRNd (bottom). In both cases electrode tracks through the NPH (red arrow) and PGRNd (blue arrow) are visible. **C, D)** Examples of representative AHV cells localized to the NPH (C) and PGRNd (D). Cells 2, 8, 12: asymmetric cells. Cells 1, 7: asymmetric-unresponsive cells. Cells 3, 4, 5, 10, 11: symmetric cells. Cells 6 and 9: inverted cells. Cells 3 and 12 are considered offset cells where the minimum firing rate occurred at ~100°/sec for cell 3 (blue arrowhead); for cell 12 the maximum firing rate occurred at ~50°/sec for the inverted cell (blue arrowhead). The location of different AHV cell types is shown for each track. There was no apparent organized localization for any of the AHV cell types. Cells 1, 2, 3, 5, 9, 10, 11 recorded from the right hemisphere; cells 4, 6, 7, 8, 12 recorded from the left hemisphere. Labels for all plots, as well as CW and CCW values, are as depicted for cell 10. Abbreviations: asc7: genu of the facial nerve; CI: caudal interstitial nucleus of the medial longitudinal fasciculus; g7: genu of the facial nerve; mlf: medial longitudinal fasciculus; MVN: medial vestibular nucleus; NPH: nucleus prepositus hypoglossi; PGRNd: paragigantocellularis reticular nucleus dorsal; V4: 4^th^ ventricle.

**Figure 3.**
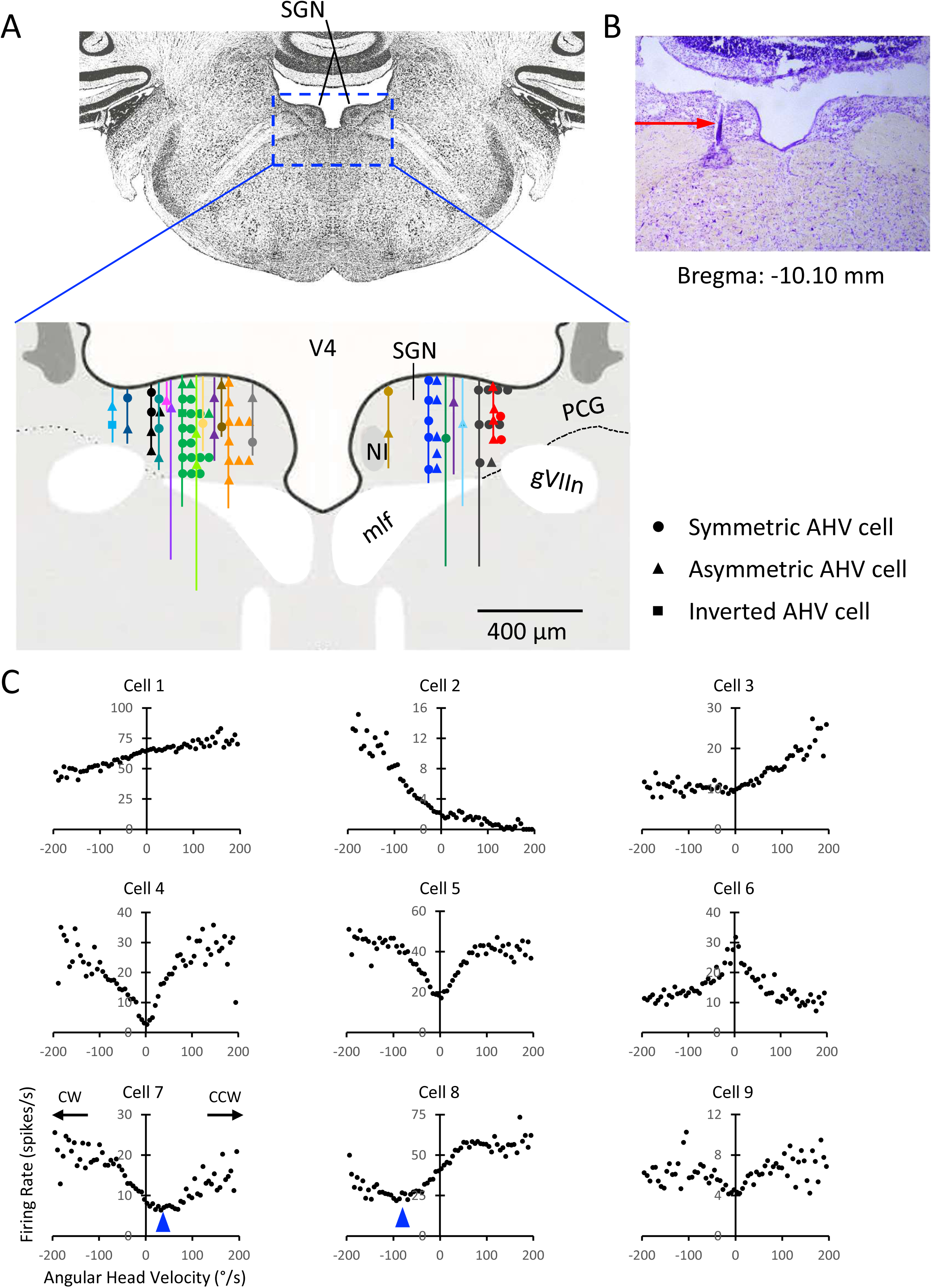
SGN AHV cells. **A)** Photograph of coronal Nissl-stained section at about −10.30 posterior to bregma. Magnified view is a schematic illustration showing the location of AHV cells along with the electrode tracks from all rats passing through the SGN. Each color track represents a different animal. **B)** A representative thionin-stained section showing an electrode track (red arrow) passing through the SGN. **C)** Examples of representative AHV cells. Cells 1-3: asymmetric cells; cells 4-5: symmetric cells; cell 6: inverted cell; cells 7-8 show examples of offset cells where the minimum firing rate is not at 0 °/sec (blue arrowheads). Cell 9 is a symmetric AHV cell that marginally passed the correlation, slope, and shuffle criteria for AHV, but was not identified by the GLM analyses as an AHV cell. The location of different AHV cell types is shown for each track. There was no apparent organized localization for any of the AHV cell types. Cells 2, 5, 8, 9 recorded from the right hemisphere; cells 1, 3, 4, 6, 7 recorded from the left hemisphere. CW and CCW values and labels for all plots are as shown for cell 7. Abbreviations as in Fig. 2 plus NI: Nucleus incertus; PCG: pontine central gray; SGN: supragenual nucleus.

Of the cells in the dataset, 231 cells (44.2%) contained tuning curves that clearly looked AHV sensitive. All except five of these cells passed our classification criteria for AHV modulation, which required: 1) the tuning curve containing an absolute value correlation ≥ 0.5 and an absolute value slope ≥ 0.025 spikes/s for at least one turn direction (CW or CCW) (henceforth referred to as threshold values). Most cells that passed our criteria threshold also passed the 95^th^ percentile shuffle test (222/231; 96.1%) and indeed most of them passed the 99^th^ percentile (211/231; 91.3%). Of the five cells that did not pass the criteria threshold, two of them passed the 99^th^ percentile shuffle test, one cell passed the 95^th^ percentile, and the tuning curves for the remaining two cells clearly looked AHV sensitive. These five cells were therefore included in the analyses and their AHV tuning curves are shown in Extended Data Figure S1. Based on this dataset, the number of cells tuned to AHV were 37.1% for NPH, 51.3% for SGN, and 44.9% for PGRNd. Many of the firing rate × AHV functions had good linear fits with steep slopes at low AHVs (0 to ± 90°/s), but at high AHVs the linear relationships became less striking, and the slopes became less steep. These characteristics were particularly evident for symmetric AHV cells (see below).

### AHV cell general properties

The AHV tuning curves could be divided into two major types depending on whether cell firing increased with increasing AHV in one (CW or CCW) or both turn directions; these two broad categories were referred to as asymmetric and symmetric AHV cells, respectively. Figures 2C-D show representative examples of AHV cells from NPH and PGRNd, respectively, and Figure 3C show representative examples from SGN. Symmetric AHV cells had generally V-shaped, bilaterally symmetric tuning curves with roughly equal magnitude responses but opposite sign for CW and CCW directions. In addition, symmetric cells could be further classified based on whether cell firing rates increased (Fig. 2C-D, cells 3-5, 10-11; Fig. 3C, cells 4-5, 7, 9) or decreased with increasing AHV (Fig. 2C-D, cells 6, 9; Fig. 3C, cell 6). Cells that displayed decreasing firing rates with increasing AHVs were referred to as ‘inverted AHV cells’. Most symmetric AHV cells had tuning curves where the significant correlations were only present between ± 0-90°/s; values ≥ 90°/s and ≤ −90°/s were either flat (slope = 0 spikes/°/s) or rather quite variable (e.g., see Fig. 2C-D, cells 5 and 11; Fig. 3C, cells 5 and 9). For asymmetric cells, firing rates increased linearly with AHV in only one turn direction, but could be further subdivided into two categories based on their response in the opposite turn direction. Some asymmetric AHV cells had decreasing firing rates in the opposite direction (referred to simply as asymmetric cells) (Fig. 2C-D, cells 2, 8, 12; Fig. 3C, cells 1, 8), while other asymmetric cells were not modulated by AHV in the opposite direction (referred to as asymmetric-unresponsive) (Fig. 2C-D, cells 1, 7; Fig. 3C, cells 2-3). Each AHV cell type was observed in all three brain areas and is described in more detail below. Over all AHV cells, we found that the AHV tuning curve slope correlated well with the cell’s baseline firing rate (Pearson *r* = 0.37, p =2.01 × 10^−17^), but correlated only weakly with the Pearson *r* (*r* =0.1, p = 0.03).

In order to better characterize the diversity of AHV tuning observed, we developed a symmetry measure (normalized turn bias, see Methods), which compared the differences between CW and CCW slopes to that of a perfectly symmetric cell with both slopes equal to the cell’s maximum slope (CW or CCW). Thus, a perfectly symmetric cell will have a normalized turn bias near 0. Plots were constructed that plotted the CW vs. CCW slope values across all AHV cells for each brain area (Figs. 4, 5, 7). For each plot, symmetric cells are located along the negative unity line in quadrants 2 and 4 and purely asymmetric cells lie along the unity line in Quadrants 1 and 3. Asymmetric cells that are neither purely symmetric or asymmetric cluster around the x and y axes. For classification purposes, we defined symmetric cells as having a normalized turn bias < 0.3 and symmetric cells as having a normalized turn bias > 0.7, while cells with a symmetric turn bias between 0.3 - 0.7 were classified as asymmetric-unresponsive.

**Figure 4.**
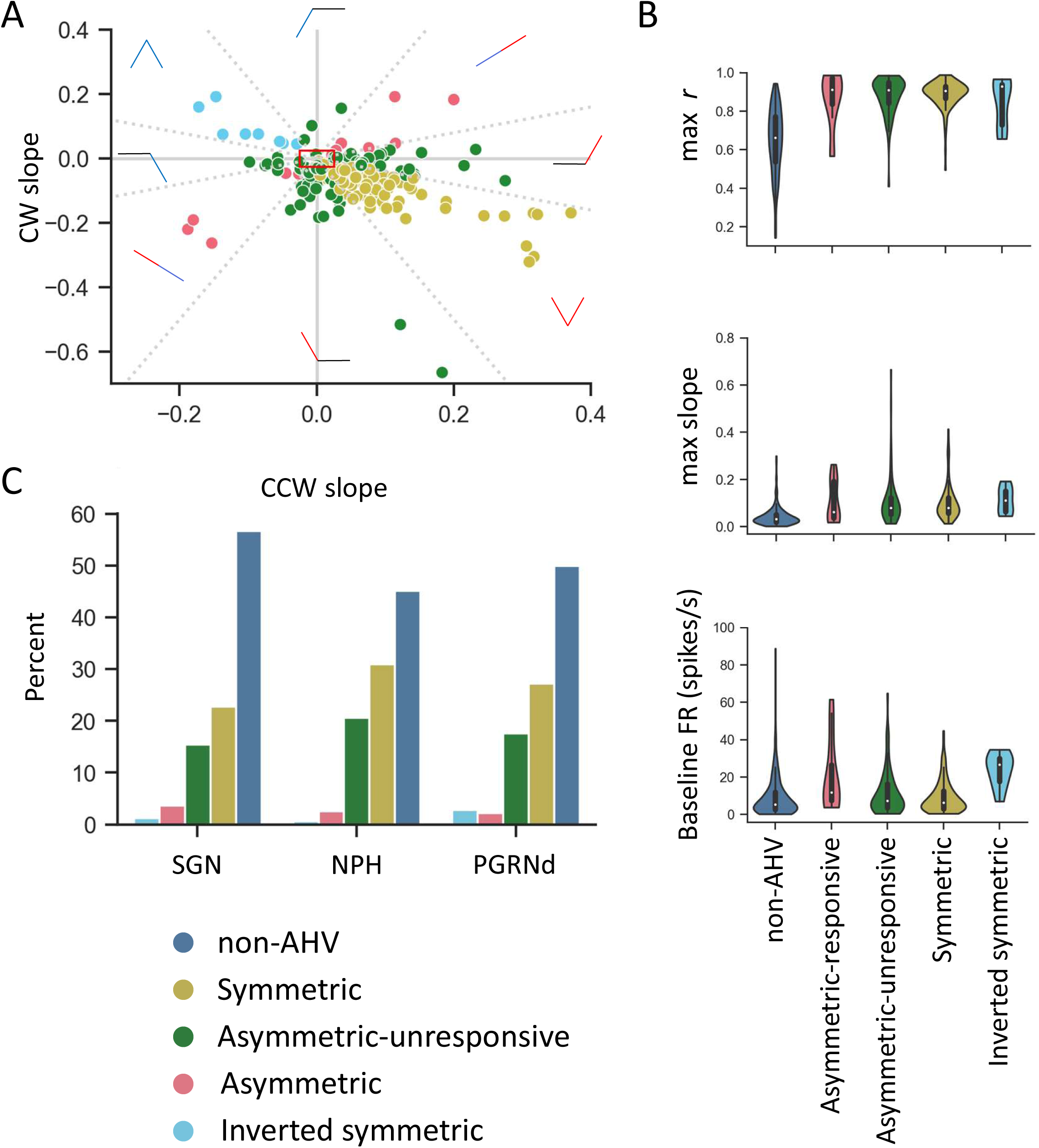
**A)** A scatterplot of the magnitude of the slopes for all AHV cells recorded, color-coded based on cell type. Dashed gray lines indicate the cutoffs dividing AHV cell types. The red box indicates the boundaries of the threshold slope criterion (absolute value of slope ≥ 0.025). Each point represents the CW and CCW slope of an AHV cell. Cells are color-coded to represent different AHV cell types. The expected AHV tuning curve shape for each AHV cell type is shown around the perimeter within the region of the plot containing that cell type. Red AHV icons indicate portions of the tuning curves that show a positive relationship between increasing AHV and firing rate, blue lines indicate a negative relationship, and black lines indicate no relationship. **B)** AHV tuning curve measures compared across different AHV cell types and non-AHV cells. **C)** Proportions of each AHV cell type in each brain region. There were no significant differences between the three brain areas.

Using this method, the CW and CCW slopes for each cell are shown in Figure 4A with colors indicating their classification. The 8 zones used for symmetry classification (two each for symmetric and asymmetric and four for asymmetric-unresponsive) are well described by the lines y= ±0.4x and y= ±(1/0.4)x shown as dotted gray lines on the plot. For comparing the strength of AHV tuning across different brain areas or hemispheres, we used the absolute value of the maximum slope and Pearson correlation that passed our selection criteria. If both CW and CCW turns passed our selection criteria, we used the maximum slope and correlation value between the CW and CCW turns. We note that a few AHV cells did not fit well into this classification scheme. In particular, there was a small number of cells, referred to as ‘offset’ AHV cells that had their firing rate minimum (or maximum in the case of inverted cells) not centered on 0°/s, but rather centered on a value that was shifted 5-10°/sec CW or CCW from 0 (see blue arrowheads in Fig. 2C, cells 3, 6, 12; Fig. 3C, cells 7, 8). Offset cells formed a very small minority of the total AHV cells in the recorded regions with two offset cells recorded in NPH and one cell in PGRNd.

Not all cells classified as AHV had robust AHV sensitivity, although they still passed our threshold criteria. These cells had *r* and slope values that only marginally passed the threshold criteria; an example of such a cell is cell 9 in Fig. 3C. This symmetric AHV cell from the SGN had values in the ± 90°/s range, which qualified it for classifying it as an AHV cell: 0-90 CW *r*: −0.930, slope: −0.035; 0-90 CCW *r*: 0.819, slope: 0.037. The cell also passed the shuffle procedure and the response was stable across the recording session; the correlation (*r*) between the first and second halves of the session between −90 and +90 °/s was 0.616.

Two-way ANOVAs of AHV tuning curve characteristics (correlation, slope, baseline firing rate) showed that the maximum Pearson correlation (max *r*) did not differ between the AHV cell types or brain regions (*p* > 0.05 in both cases; Fig. 4B top). Maximum slope also did not differ between AHV cell types or brain regions (*p* > 0.05 in both cases), but did exhibit a weak interaction (F_(4,451)_ = 3.67, p=0.006; Fig. 4B middle). Post-hoc tests indicated that this interaction was because PGRNd differed from the other two brain regions, with symmetric cells exhibiting increased slopes and asymmetric-unresponsive cells exhibiting decreased slopes. Baseline firing rates did not differ between the AHV cell types (*p* > 0.05), but did differ significantly across brain regions (F_(2,451)_ = 16.63, p < 10^−6^). Post-hoc tests again indicated that this effect was because PGRNd differed from the other two brain regions by exhibiting significantly higher baseline firing rates (PGRNd vs SGN & NPH: *p* < 10^−4^, *p* = .71 for SGN vs NPH; Fig. 4B middle).

The proportion of each AHV cell type by brain area is shown in Figure 4C. There was no prominent difference in the proportions of cell type across the three brain areas (χ^2^ _(4, 451)_, p=0.22). In addition, the number of inverted AHV cells we identified in each brain area was low, with 2 cells in NPH, 1 cell in SGN, and 5 cells in PGRNd. Figures 2D (cell 9), 3C (cell 6), and 8A (third cell from left) show example AHV inverted cells. The proportion of symmetric vs. asymmetric cell types did not differ by brain area (χ^2^ _(4, 451)_, p=0.22).

To develop biologically relevant models of HD function, it is important to identify not only the physiological cell types but also to understand whether the connections are excitatory or inhibitory onto the downstream HD network. In more cortical regions, putative inhibitory cells can be distinguished from putative pyramidal cells by a combination of firing rate and spike width values (McCormick et al., 1985; Bartho et al., 2004). For example, cells with fast firing rates and short spike widths are thought to be inhibitory. We therefore examined whether there were any differences in spike width between different AHV cell types and across the three different brain areas. In brief, we did not find any differences in spike width (see Extended Data Figure S2).

### Cell firing stability

To insure that AHV sensitivity was stable across the entire recording session, we compared correlation and slope measures between the first and second halves of a recording session. We examined correlation values between the 1^st^ and 2^nd^ halves for both the CW and CCW portions of each cell’s AHV tuning curve, as well as the entire tuning curve’s range (± 0-204°/s). Overall, 94.8% of the 231 AHV cells had correlations ≥ 0.5 for either the CW or CCW portion of its tuning curve. For NPH, 60 out of 66 AHV cells had correlations ≥ 0.5; for SGN, 78 out of 81 AHV cells had correlations ≥ 0.5; for PGRNd, 81 out of 84 AHV cells had correlations ≥ 0.5. For the remaining 12 cells that had correlations < 0.5, all except 2 cells (in NPH) had correlations ≥ 0.4. The mean 1^st^ vs. 2^nd^ half correlations for the portion of the cell’s tuning curve that was ≥ 0.5 were NPH: 0.694 ± 019 (median: 0.692), SGN: 0.742 ± 0.016 (median: 0.760), and PGRNd: 0.696 ± 0.014 (median: 0.650).

### Cell firing related to ipsiversive vs. contraversive head turns

Because the rats were implanted in either the right or left hemispheres, asymmetrical AHV cells were further examined to determine whether increased firing was associated with ipsiversive or contraversive head turns in relation to which hemisphere the cell was recorded from. Figure 5A depicts a scattergram of the best-fit slopes for CW vs. CCW head turns for all asymmetric and asymmetric-unresponsive cells based on whether the turn was ipsiversive (turning towards the hemisphere the cell was recorded from) or contraversive (turning away from the hemisphere the cell was recorded from). Figure 5B displays the percentage of asymmetric cells showing ipsiversive and contraversive firing across the three brain areas. Although there was a trend for ipsiversive head turns leading to increased firing in SGN, this trend was not significant. A Chi-square test for independence showed that the proportion of cells selective for contraversive turns was not different between brain areas (χ^2^ _(2,97)_ = 0.68, p = 0.42). Thus, both ipsiversive and contraversive cells could be found within the same hemisphere. Figure 5C shows two asymmetric AHV cells that were recorded simultaneously from the same hemisphere (left) in NPH; the two recordings occurred on the same day from different electrodes. The firing rate increases in the CW direction for the cell on the left (contraversive) and in the CCW direction in the cell on the right (ipsiversive). Figure 5D shows AHV tuning curve measures for the same cells as in Figure 5A compared between ipsiversive and contraversive turns for each brain area. There were no significant differences for max slope or max *r*, but there was a main effect for baseline firing rate (F_(1,91)_ = 9.55, p = 0.003, interaction and main effect of brain area: p> 0.19). Tukey post-hoc tests showed significant baseline firing rate differences between contraversive and ipsiversive selective cells (p = 0.001).

**Figure 5.**
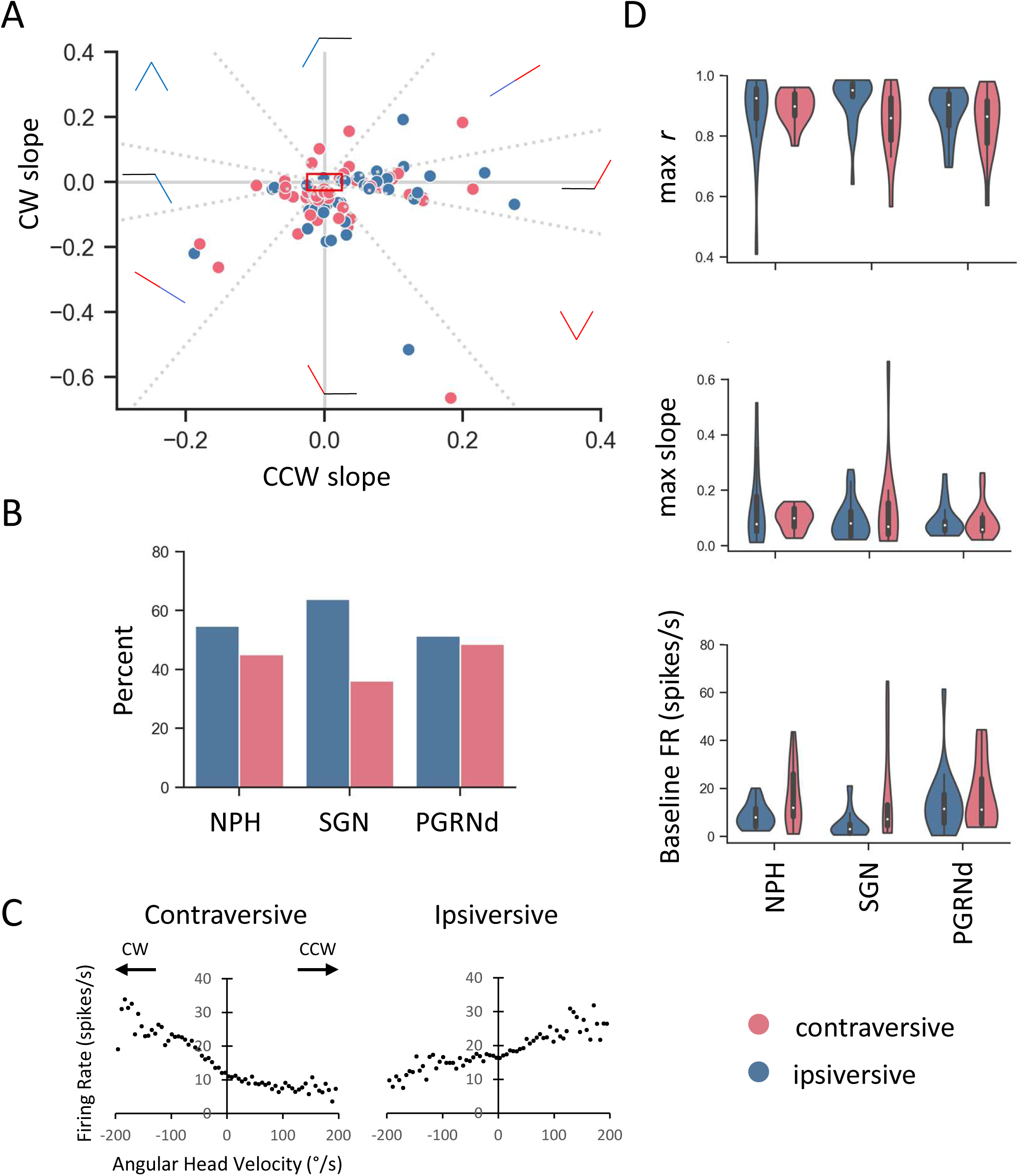
**A)** A scatterplot of the magnitude of the slopes (as in Figure 4B) showing all AHV cells with a contraversive or ipsiversive turn bias. The red box indicates the boundaries of the threshold slope criterion (absolute value of slope ≥ 0.025). The expected AHV tuning curve shape for each AHV cell type is shown around the perimeter within the region of the plot containing that cell type. Red AHV icons indicate portions of the tuning curves that show a positive relationship between increasing AHV and firing rate, blue lines indicate a negative relationship, and black lines indicate no relationship. **B)** Proportions of contraversive and ipsiversive AHV cells in each brain area. **C)** Two asymmetric AHV cells recorded simultaneously on different electrodes that were in close proximity to one another in the left NPH. Note that the firing rate increases in the CW direction for one cell (left, contraversive) and in the CCW direction in the other cell (right, ipsiversive). **D)** AHV tuning curve measures (max *r*, max slope, baseline firing rate) compared between ipsiversive and contraversive cells for each brain area.

### Many AHV cells also respond to linear velocity

Previous studies have reported that HD cells and AHV cells in the lateral mammillary nucleus and DTN correlated to linear velocity (Stackman and Taube, 1998; Bassett et al., 2001). We therefore analyzed linear velocity tuning curves for cells in NPH, SGN, and PGRNd. Cells were classified as modulated by linear velocity if 1) if the best-fit line of their firing rate vs. linear velocity tuning curve had a correlation > 0.7 and slope ≥ 0.1 spikes/cm/s and 2) the correlation and slope values passed a 95% shuffle procedure.

Based on this definition, 28% of all recorded cells were tuned to linear velocity (NPH: 18%, SGN: 29%, PGRNd: 35%). We found that 46% of all AHV cells across all three brain areas were modulated by linear velocity with PGRNd having the largest proportion of AHV cells that passed our criteria for linear velocity (NPH: 38%, SGN: 44%, PGRNd: 55%). Figure 6A shows 5 AHV cells that were also tuned to linear velocity. Most cells that passed the linear velocity criteria had positive slopes for their tuning curves (Fig. 6A, cells 1, 2, 5). However, similar to inverted AHV cells, we identified a few cells that had negative slopes in their linear velocity tuning curve indicating they had a decreasing firing rate as a function of increasing linear velocity (Fig. 6A, cells 3, 4). Four out of 5 inverted AHV cells from PGRNd contained a negative slope in their linear velocity tuning curve, and across all recorded cells (both AHV and non-AHV), almost all cells with negative linear velocity/firing rate correlations were found in PGRNd (14/16). AHV cells that were tuned to linear velocity formed a heterogeneous population of AHV cell types in the sense that they could be symmetric (Fig. 6A, cell 2), asymmetric (Fig. 6A, cell 4), asymmetric non-responsive (Fig. 6A, cells 1, 5), or inverted (Fig. 6A, cell 3).

**Figure 6.**
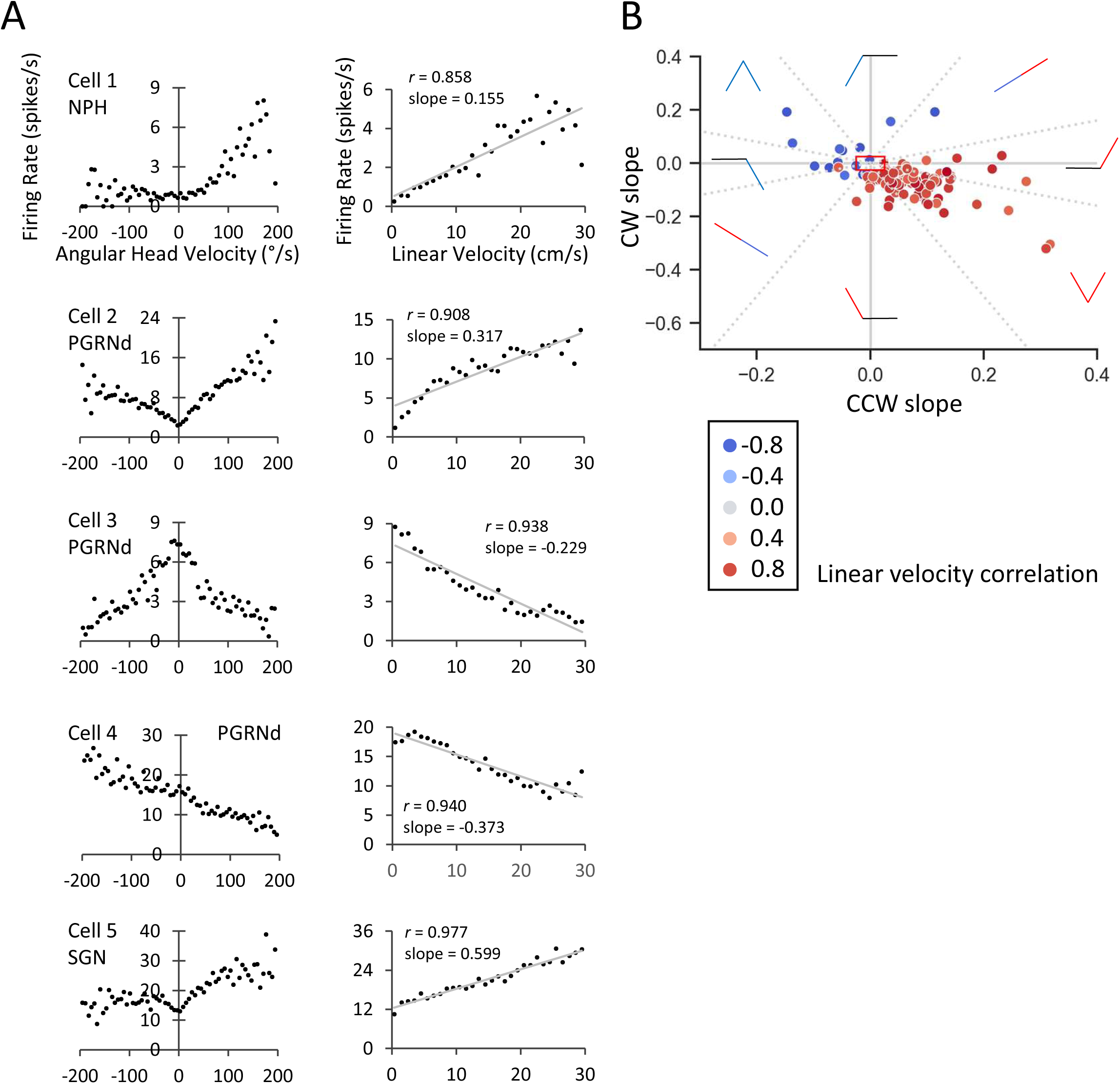
**A)** Example AHV (left) and linear velocity (right) tuning curves for 5 cells with significant tuning for both variables. **B)** A scatterplot of CW and CCW slopes (as in Figs. 4 and 5) of cells that passed threshold criteria and shuffles for both AHV and linear velocity. The red box indicates the boundaries of the threshold criterion (*r* ≥ 0.5; absolute value of slope ≥ 0.025). Red lines indicate a positive linear velocity correlation, blue lines a negative linear velocity correlation, and black lines no relationship.

On closer inspection of the relationship between linear and angular velocity values, the sign of the linear velocity slope (positive vs. negative) tended to be in the same direction as the AHV slopes (i.e., cells that coded for both AHV and linear velocity had firing rates positively correlated to angular and linear velocity, while cells that contained negative slopes for their linear velocity tuning curves had firing rate decreases in their AHV tuning curve (i.e., were inverted AHV cells). However, we identified a few (n=12) cells that contained a negative slope in their linear velocity tuning curve that were not inverted AHV cells. These cells were mostly asymmetric-unresponsive (n=10) or asymmetric (n=2)(Fig. 6A, cell 4). Figure 6B shows the same scatterplot as in Figures 4 and 5 but with the color hue indicating the linear velocity Pearson correlation. Asymmetric-unresponsive cells tend towards positive linear velocity correlations if the firing rate increases with the magnitude of AHV for either CW or CCW directions. Conversely, asymmetric-unresponsive cells that decrease their firing rate with increases in AHV magnitude (i.e., ‘asymmetric-inverted’ AHV cells) also have strong negative linear velocity correlations (i.e., firing rate decreases with increasing linear velocity). Of note, and in agreement with recordings from AHV cells in the DTN and LMN (Stackman and Taube, 1998; Bassett et al., 2001), cells with positive firing rate × linear velocity correlations make up the majority of the population of cells with significant AHV and linear velocity tuning (~85%).

Because linear and angular head movements can be difficult to distinguish based on the tracking of two points, we applied an additional analysis using a GLM to determine whether cells coded for linear velocity, AHV, or both (see Methods). Figure 7A displays GLM classification type as a proportion of the total population for all cells divided by the number of variables conjunctively classified (e.g., cells that only code for 0-1 variables are on the left while cells that code for all possible variables in the model are on the right). The same information divided by brain area is shown on the right. For all brain regions, the largest number of cells was classified as linear velocity × AHV, followed by AHV alone, and then HD alone. As expected, the GLM and threshold criterion methods for classifying AHV cells overlapped considerably, although not perfectly. 72% of the cells that passed the threshold and shuffle criteria were classified as AHV by the GLM, while 83% of the cells chosen by the GLM as AHV passed the threshold/shuffle criteria. In contrast, there was less agreement between the GLM classification for linear velocity tuning and our method of selecting linear cells based on threshold criteria (correlation and slope) and shuffle analyses (see Methods). Here, only 52% of the cells that were classified as linear velocity-tuned by the GLM also passed the threshold criteria and the shuffle procedure. Similarly, only 56% of the cells that passed the linear threshold and shuffle analyses were classified as linear velocity tuned by the GLM. We further noticed that the GLM tended to classify asymmetric and asymmetric-unresponsive cells as solely tuned to AHV, while symmetric AHV cells were classified as conjunctively coding AHV and linear velocity. Cells classified as both AHV and linear velocity-tuned by the GLM had more symmetric AHV tuning curves than cells that were classified as encoding AHV alone as evidenced by smaller normalized turn bias values (t(87) = 6.43, p =6.68*10^−9^) (Fig. 7B).

**Figure 7.**
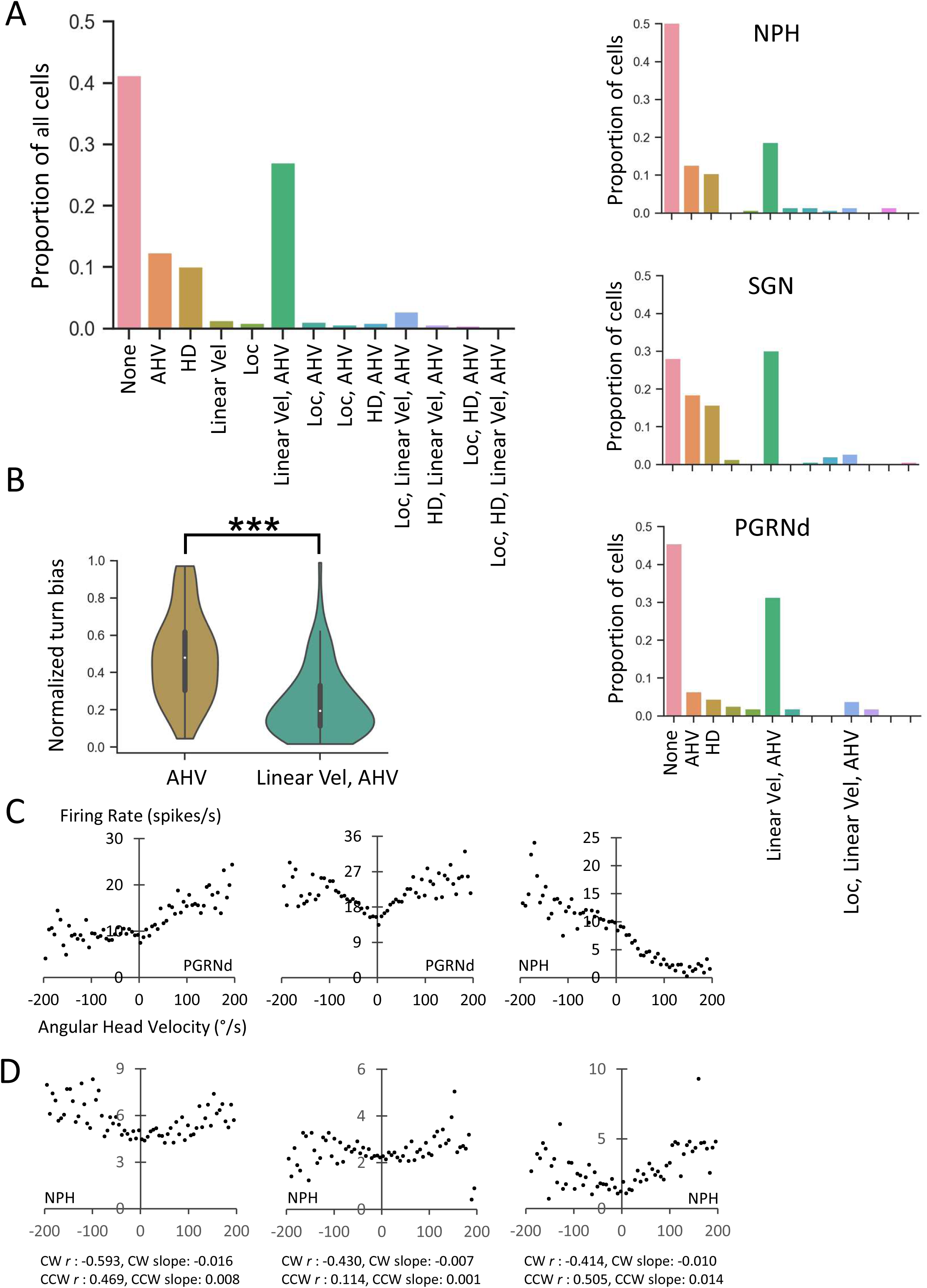
**A) Left:** Proportions of GLM classifications using head direction (HD), AHV, linear velocity, and location (Loc) as variables. **Right:** Same as **left**, but for each brain area. **B)** Bar graph comparing symmetry (normalized turn bias) in cells that the GLM classified as only AHV or conjunctively coding AHV and linear velocity. **C)** Examples of three AHV cells that passed the threshold criteria for correlation, slope, and shuffle procedure, but did not pass the GLM criteria for AHV. **D)** Examples of three non-AHV cells that passed the GLM criteria for AHV, but did not pass the threshold criteria. Note that in each case either the cell’s CW/CCW correlation *r* or slope values did not reach the threshold criteria for categorizing the cell as an AHV cell.

While all brain regions have similar proportions of AHV symmetry cell types, PRGNd had a larger proportion of AHV × linear velocity conjunctive cells and a lower proportion of cells classified by the GLM as AHV or HD alone than NPH or SGN (χ^2^ _(4,219)_ = 14.49, p = 0.006)). Post-hoc Chi-square tests removing one area at a time show significance only when PGRNd is included in the comparison (NPH excluded (χ^2^ _(1,154)_ = 10.23, p < 0.006; SGN excluded (χ^2^ _(1,114)_ = 12.09, p = 0.002; PGRNd excluded (χ^2^ _(1,144)_ = 0.57, p=0.75). Regardless of whether the cells can be defined as tuned for linear velocity or AHV, it is clear that the three brain regions show a remarkable similarity in the information they represent. Nonetheless, it is important to note that there were a large number of cells that appeared tuned to AHV that were not classified as such by the GLM (Fig. 7C; also see Fig. 2, cell 4 and Fig. 3, cell 6 for other examples). Conversely, the GLM classified a number of cells whose firing was modulated by AHV that were not reflected in their tuning curves (Fig. 7D). This result was also true for linear velocity. Fig. 8A shows two cells that were tuned to linear velocity based on the threshold criteria, but were not selected by the GLM as linear velocity tuned. Conversely, Fig. 8B shows two cells that the GLM selected as tuned to LV, but were not classified as such based on the threshold criterion and their linear velocity tuning curves were relatively flat and did not look particularly sensitive to LV. These results provide an important cautionary note for not relying entirely on a GLM analysis to demonstrate linear or AHV firing rate sensitivity.

**Figure 8.**
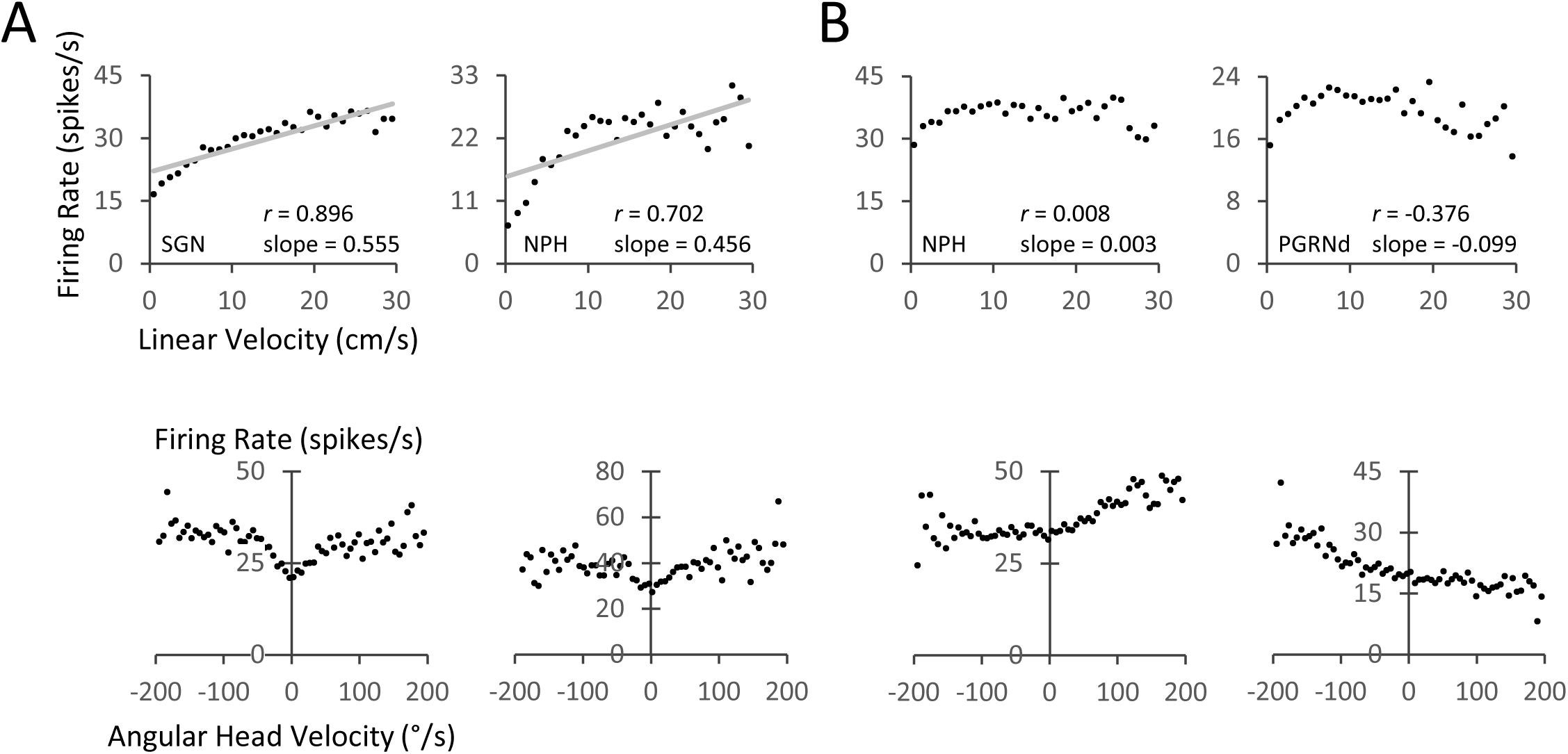
Linear Velocity and GLM. **A)** Linear velocity tuning curves for two representative cells that passed the linear velocity threshold criteria (correlation, slope, and shuffle), but were not selected by the GLM analysis as linear velocity sensitive. **B)** Linear velocity tuning curves for two representative cells that did not pass the linear velocity threshold criteria (correlation, slope, and shuffle), but were selected by the GLM analysis as linear velocity sensitive. The corresponding AHV tuning curves are shown below each cell. The cell on the right in *A* did not meet threshold criteria to be classified as an AHV cell.

### Head direction modulation

In terms of the HD cell system, the most upstream (afferent) recordings of HD cells were reported in the DTN (Sharp et al., 2001b). All three brain areas recorded here are afferent to DTN, but situated downstream from the vestibular nuclei. The GLM classified 54 cells (AHV + non-AHV; 10.9% of total recorded cells) from the NPH, SGN, and PGRNd as modulated by HD. However, except for one cell (see below), the cells classified as directionally sensitive by the GLM did not appear to be modulated by HD based on their firing rate vs. HD tuning curves (Fig. 9A). Indeed, their tuning curves did not resemble classic, let alone HD-modulated, tuning curves from other brain areas (e.g., see Fig. 6 of Bassett and Taube, 2001; Fig. 1C of Clark et al., 2023). Median half-session stability (Pearson’s *r* between 1^st^ and 2^nd^ halves) for these cells was 0.15 and the median Rayleigh vector length was 0.09 – both exceptionally low values for ‘normal’ HD cells. Figure 9E shows a scatterplot of the distribution of Rayleigh vector length values and half-session stability. Note that almost all the cells have a half session stability < 0.5 and a tuning strength (Rayleigh r) < 0.3. The one HD-modulated cell discussed below is highlighted by a square symbol. Taken together with the questionable effectiveness of the GLM analyses for selecting for linear and angular head velocity sensitivities (see above), the use of GLM methods for classifying cells as HD cells in subcortical structures is not very reliable.

**Figure 9.**
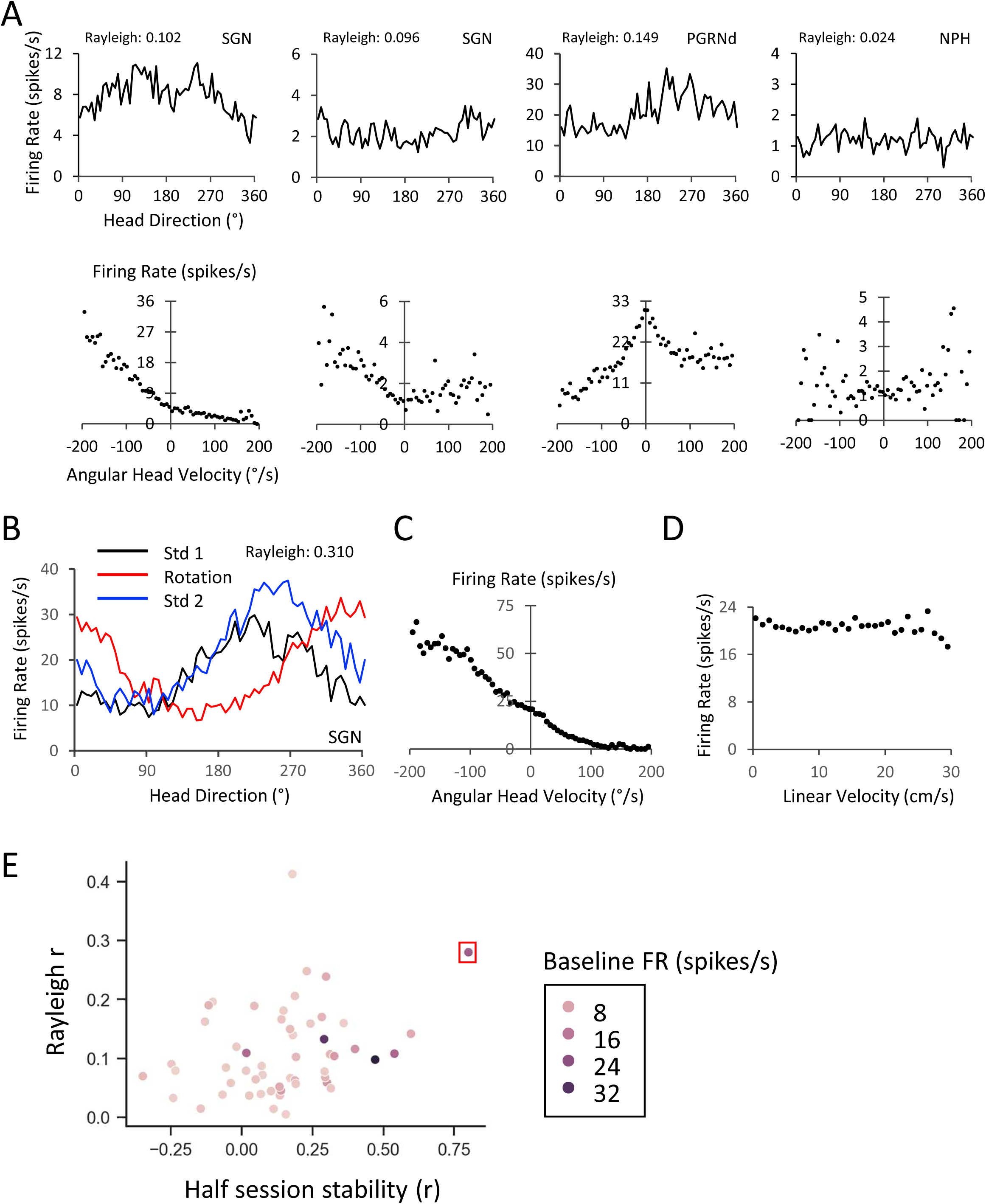
**A)** Examples of three AHV cells and one non-AHV cell (far right) that passed the HD GLM criteria, but did not have significant directional tuning by classic classification criteria (Rayleigh r > 0.4, directional firing ranges < 120°, peak firing rates > 5 spikes/s). HD vs. firing rate plots are shown in the top row and their corresponding AHV tuning plots are shown in the bottom row. **B)** Head direction tuning curves for the one SGN cell identified as HD modulated during two standard conditions (black and blue lines) and an intervening session, where the prominent visual cue in the recording arena was rotated by 90°. **C)** The HD cell in *B* was also significantly tuned to AHV with an asymmetric tuning curve. **D)** The same cell’s tuning curve for linear velocity, showing no firing rate modulation with changes in linear velocity. **E)** Summary plot of all cells identified as ‘HD’ by the GLM. Almost all of the cells have a half session stability < 0.5 and a tuning strength (Rayleigh r) < 0.3. The HD-modulated cell in *B* is highlighted by a red square symbol.

Only one asymmetric AHV cell in SGN had a tuning curve that was recognizable as an HD cell. Figure 9B-D displays the firing rate by HD for this cell as well as its AHV tuning curve; it did not show any linear velocity sensitivity. The cell’s firing for both HD and AHV was stable across the 16 min session. Half session correlations (Pearson’s *r*) for AHV and HD were: 0.955 and 0.798, respectively. Unlike brain regions that contain a large proportion of HD cells, such as the anterior dorsal thalamus and postsubiculum, the directional firing range (tuning width) of the SGN cell (259.1°) was much wider, and more closely resembled those from HD cells that are better characterized as HD-modulated cells within the DTN (Bassett and Taube, 2001) or conjunctive HD cells in the postrhinal cortex (LaChance et al., 2022). Among the cells classified as HD by the GLM, this cell had the highest half-session stability and a modest Rayleigh r (0.310).

We also conducted a cue rotation and a cue removal session on this cell. As with normal HD cells from other brain areas, rotation of a prominent cue card (landmark) attached to the inside wall of the cylinder led to a corresponding shift of equal magnitude in the cell’s preferred firing direction (Fig. 9B, red line). The cell’s preferred firing direction also remained stable in the absence of visual cues in a session conducted in the dark with the cue card removed, which is similar to that seen for HD cells in other brain areas (Taube et al., 1990; Taube, 1995). Although a previous study reported that visual spatial information gains control of the HD signal at the level of the LMN by way of direct projections from the postsubiculum (Yoder et al., 2015), the finding that this SGN cell responded to rotation of the visual landmark cue indicates that visual landmark information is capable of being conveyed to the SGN at early stages of the HD circuitry - prior to the inputs into LMN. It is intriguing to consider how this visual information reaches the SGN. Although the SGN receives direct input from the vestibular nuclei, Nishilke et al. (2000) in examining cortical projections to the vestibular nuclei, did not identify the visual cortex as among the areas projecting there. The SGN, however, is reciprocally connected with the DTN (Biazoli et al., 2006) and the DTN receives projections from both the LMN (Hayakawa and Zyo, 1990) and retrosplenial cortex (Mehlman et al., 2021), an area known to play a role in processing landmark information (Clark et al., 2010; Jacob et al., 2017). Therefore, it’s possible that visual landmark information is conveyed to the SGN via these feedback pathways.

### Brainstem AHV cells show mixed AHV tuning responses during passive rotation

Previous studies in head-unrestrained monkeys have shown that the firing of vestibular nuclei neurons are markedly suppressed (~70%) during actively generated head turns compared to passive turns of the head (Roy and Cullen 2001, 2004). These findings have pointed to a conundrum: how can HD cells generate their signal if the upstream vestibular signal is reduced markedly during an active head turn (Shinder and Taube, 2014; Cullen and Taube, 2017)? In addition, Sharp et al. (2001b) recorded from single neurons in the DTN under both active and passive (hand-held) conditions and reported a variety of responses across different cells. The AHV sensitivity of some DTN cells was disrupted during passive head turns, while other DTN cells maintained their AHV sensitivity. Because NPH, SGN, and PGRNd neurons are situated in between the vestibular nuclei and the DTN, and that the NPH and SGN both project directly to DTN, it was important to determine how AHV cells in these brainstem areas respond during passive head turns compared to active head turns. We used two approaches across each brain area: 1) we monitored some AHV cells while the rat was hand-held and lightly restrained while it was passively rotated back-and-forth in about 90° arcs sampling the entire 360° range of possible HDs, 2) we monitored other AHV cells while the rat was head-fixed in a small cylinder that sat atop a turntable that could be rotated freely about the vertical axis (see Fig. 1 in Shinder and Taube, 2011).

For both approaches, AHV cells were first recorded for 16 min in a freely-moving session, followed by a 1 min session, where the rat was firmly wrapped in a cloth towel (hand-held session) or placed into a restraint device (head-fixed session) and then rotated passively back-and-forth through a range of 180° at various speeds (0-270°/sec). Following the passive session, the cells were monitored a second time in a freely-moving session. To be included in the analyses below, cells had to have an AHV tuning curve that correlated well to the first active session’s tuning curve (Pearson’s *r* ≥ 0.50). In some cases (n=5) the second active session was conducted on the next 1-2 days following the passive session. 18 passive sessions were from hand-held sessions (5 NPH cells, 5 SGN cells, 8 PGRNd cells) and 9 passive sessions were from head-fixed sessions (2 NPH cells, 2 SGN cells, 5 PGRNd cells).

Of the 27 AHV cells tested, 8 cells were classified with symmetric tuning curves (NPH=2, SGN=0, PGRNd=4, 2 inverted) and 19 cells were classified with asymmetric tuning curves (NPH=5, SGN=7, PGRNd=7). Three SGN cells were recorded in multiple passive sessions across different days. In each case the cell’s response during the first passive session was the same during subsequent passive sessions.

Similar to the responses reported for DTN AHV cells, we also found a mixture of responses across the NPH, SGN, and PGRNd, with some cells maintaining a similar AHV sensitive tuning curve during the passive session compared to the active sessions, while other cells lost their AHV sensitivity. This mixture of responses was also true for the two methods of passively rotating the rats, as well as for the different AHV cell types. Figure 10 shows eight representative responses across the three brain areas, across different types of AHV cells, and across the two methodological approaches (head-fixed vs. hand-held). Cells in *C*, *E*, and *F* were classified as having lost their AHV-specific firing during the passive session or having it substantially attenuated. Cells in *A* and *D* were classified as maintaining their AHV-specific firing during the passive session, similar to that observed in the active sessions. Cells in *B*, *G*, and *H* were considered to have maintained their AHV-specific firing in the passive session, but having a different response pattern compared to the active sessions. The symmetric AHV cells in *B* and *G* became more asymmetric in the passive session and the inverted AHV cell in *H* changed its characteristics completely and became a symmetric AHV cell in the passive session. Note that the scale for the ordinate often changes between sessions. Nonetheless, the overall pattern of activation, in terms of the shape of the tuning curve and whether AHV-specific firing was maintained, can still be discerned from each plot. Table 1 summarizes AHV cell responses to passive head turns.

**Figure 10.**
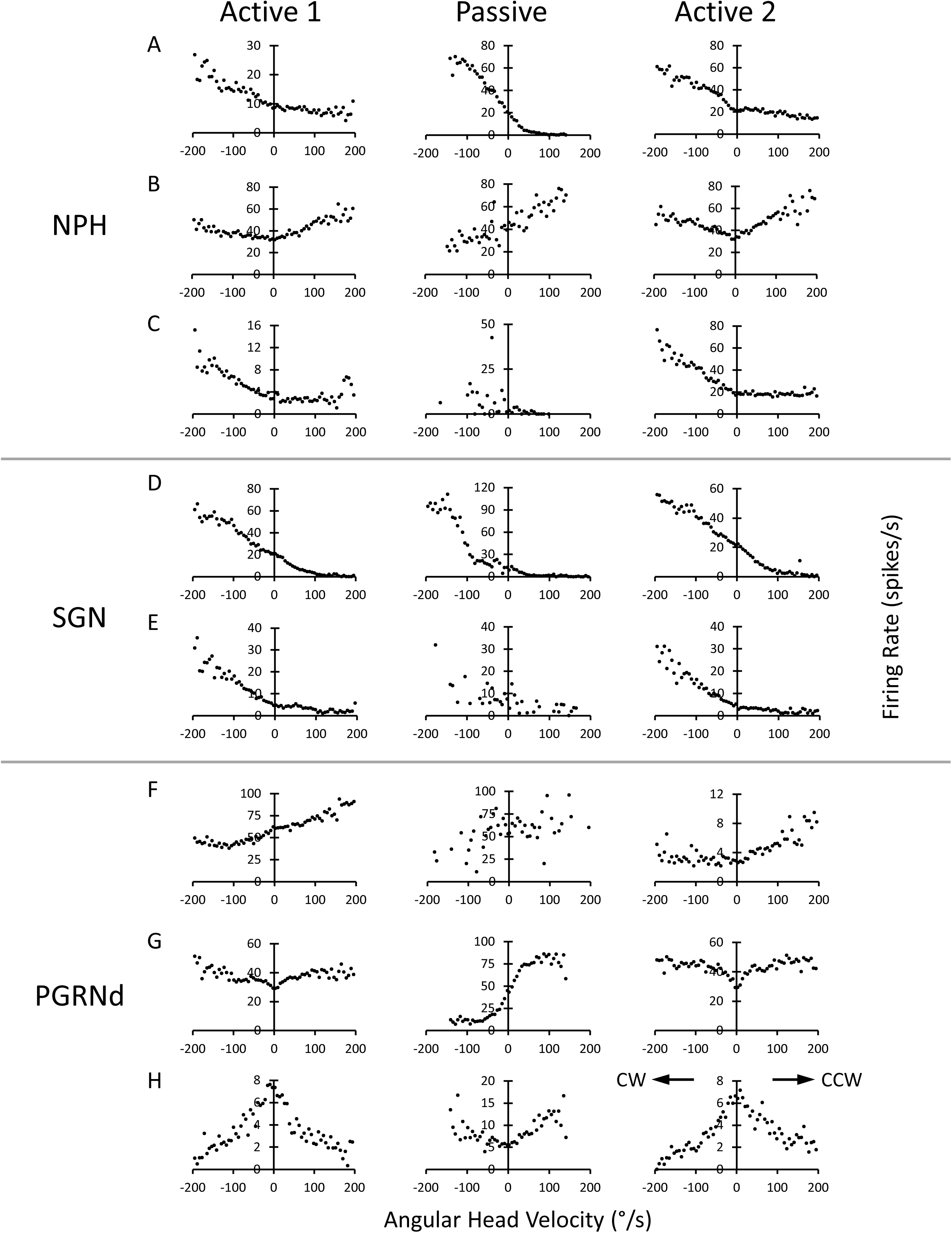
Active – Passive Responses. Tuning curves for active-passive-active sessions. **A-C)** cells from NPH, **D-E)** cells from SGN, **F-H)** cells from PGRNd. Cells in *A*, *D*, *G*, and *H* were recorded from rats which were head-fixed for the passive session; Cells in *B*, *C*, *E*, and *F* were recorded from rats which were hand-held and rotated back-and-forth in the passive session. Cells in *A* and *C-F* were classified as asymmetric AHV cells; the cells in *B* and *G* were classified as symmetric AHV cells, and the cell in *H* was classified as an inverted AHV cell. CW and CCW turns for all plots are shown in the right plot in *H*. See text for further details in terms of different response patterns. The cells in *B, C, E, F, G,* and *H* were also classified as LV-tuned cells based on threshold criteria.

**Table 1:**
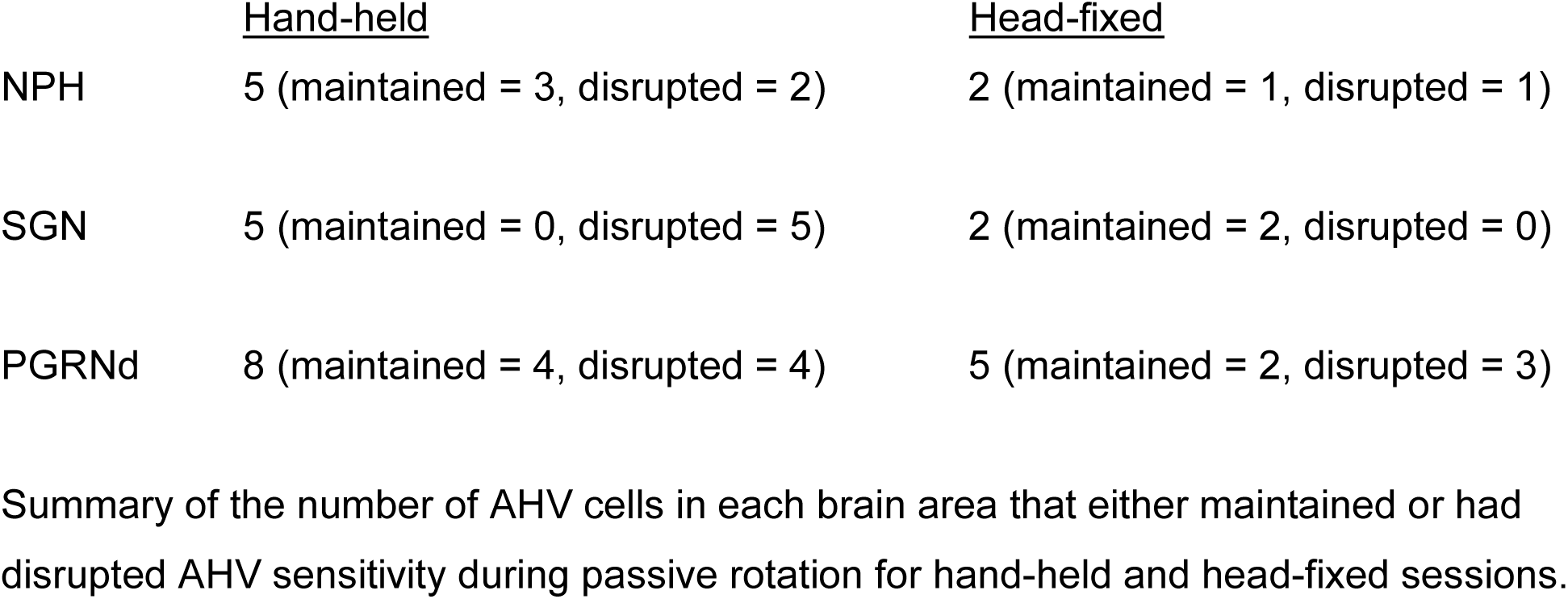
Summary of AHV cell responses to passive head turns

|  | <u>Hand-held</u> | <u>Head-fixed</u> |
| --- | --- | --- |
| NPH | 5 (maintained = 3, disrupted = 2) | 2 (maintained = 1, disrupted = 1) |
| SGN | 5 (maintained = 0, disrupted = 5) | 2 (maintained = 2, disrupted = 0) |
| PGRNd | 8 (maintained = 4, disrupted = 4) | 5 (maintained = 2, disrupted = 3) |

Summary of the number of AHV cells in each brain area that either maintained or had disrupted AHV sensitivity during passive rotation for hand-held and head-fixed sessions.

## Discussion

The aim of this study was to examine the types of spatial correlates found within three key brainstem nuclei that form part of an ascending network subserving spatial orientation in general, and directional heading information specifically. Extracellular recordings were made within NPH, SGN, and PGRNd while rats foraged freely in a cylinder, and represents the first study to record neuronal activity in these brain areas in an awake behaving rat. The major spatial correlate observed within all three nuclei were cells whose firing correlated with AHV, with some cells also showing firing sensitive to linear velocity. Indeed, about half of all recorded cells in each brain area met our classification criteria for AHV. Similar to other nodes within the ascending HD circuit, two broad categories of AHV cells were observed within each of these brainstem nuclei: symmetrical and asymmetrical AHV cells (Stackman and Taube, 1998; Bassett and Taube, 2001; Sharp et al., 2001b). As in DTN and LMN, the predominant cell types were symmetric and asymmetric-unresponsive cells with positive AHV and linear velocity correlations. The proportion of AHV symmetric to asymmetric cell type did not differ between the three brainstem regions. Similarly, there were more symmetrical and asymmetrical-unresponsive cells with firing rates that increased with AHV magnitude compared to ‘inverted’ AHV cells, which had decreasing firing rates with increasing AHVs for both CW and CCW head turns. These inverted AHV cells were found in small proportions in LMN (Stackman and Taube, 1998), but were not found in DTN (Bassett et al., 2001). For asymmetrical AHV cells, there were more cells that had a non-responsive head turn direction (either CW or CCW) compared to cells that had firing rates correlating to AHV in both directions – albeit one direction with increased firing and the other direction with decreased firing, which is more typically seen in neurons in the vestibular nuclei or VIIIth nerve afferents (Miles, 1974; Massot et al., 2012). Indeed, only 13 cells (5.6%) across all three brainstem areas showed asymmetric responses similar to vestibular nuclei neurons or their afferents. While it is true that many asymmetric non-responsive cells had firing rates that hovered around 0 spikes/s in the non-modulated direction, thus precluding them from being AHV-modulated, there were many cells that had firing rates well above zero in their non-modulated direction (e.g., Fig. 3, cell 3; Fig. 5C, left). Thus, low resting firing rates cannot be an explanation for this difference with vestibular neurons.

Comparisons of AHV properties across the NPH, SGN, and PGRNd yielded few significant differences, probably reflecting, in part, the intimate interconnections among these regions and their common afferents (Liu et al., 1984; McCrea and Horn, 1996; Biazoli et al., 2006). Neither correlation coefficients, or slopes differed across brain regions or AHV cell types indicating that the strengths of AHV modulation were similar. Similar to previous recordings from NPH in anesthetized rats, we identified both ipsiversive (Type I) cells and contraversive (Type II) cells (Lannou et al., 1984), but unlike this previous study, the proportion of ipsiversive and contraversive cells were about equal (43% contra, 57% ipsi). Furthermore, Lannou et al. (1984) reported that every cell recorded from NPH responded to horizontal sinusoidal rotation in the dark, which differs from our findings in that only 37% of NPH cells were classified as AHV sensitive.

### Comparison to other studies in non-human primates or immobilized rodents

Although this experiment is the first study to record from these regions in a freely behaving rodent, other vestibular studies have found a variety of cells types that respond differently to active and passive rotations of the head, neck, and eyes (Grantyn and Berthoz, 1987; McFarland and Fuchs, 1992; Kitama et al., 1995; Dale and Cullen, 2013). The most studied of the three nuclei, NPH, is known as the oculomotor integrator (Robinson, 1989; Fukushima et al., 1992) and is thought to be primarily involved in gaze stabilization by encoding eye movement information during the vestibulo-ocular reflex, smooth pursuit, and contributes to the generation of saccades (Lopez-Barneo et al., 1982; McFarland and Fuchs, 1992; Cullen et al., 1993; Escudero et al., 1996). In non-human primates NPH contains cells that respond to eye-position and eye-velocity signals during both active and passive head movements (Dale and Cullen, 2013). However, a smaller population of cells was responsive to both head and eye movements during passive rotations of the head (Lannou et al., 1984; McFarland and Fuchs, 1992; Cullen et al., 1993). Furthermore, some cells were shown to correlate more with head movements rather than eye movements, at least during passive rotations of the head (Delgado-Garcia et al., 1989; McFarland and Fuchs, 1992). Given that eye movements were not monitored in the present study, it remains possible that some of the correlates attributed to AHV may be related to eye movements. However, eye movements typically occur with head movements in the rat (Tempia et al, 1992; Wallace et al., 2013); therefore, cell activity in our study would be highly correlated with both AHV and eye movement. PGRNd cells may also respond to the coordination of eye and head movements (i.e., eye-neck cells) during gaze shifts and when orienting towards novel targets (Grantyn and Berthoz, 1987). A second, less common, class of observed cells was vestibular-only, which responded to passive rotations of the head (Kitama et al., 1995). Future studies that co-record cellular activity in conjunction with the animal’s eye movements in a freely-moving animal would undoubtedly clarify this issue.

Although the AHV tuning curves from SGN cells resembled those from NPH and PGRNd, we identified at least one cell that was conjointly modulated by AHV and HD. This AHV × HD cell resembled those reported within DTN (Bassett and Taube, 2001; Sharp et al., 2001b). Because of SGN’s anatomical connectivity with NPH as well as other regions, it remains possible that some SGN neurons are also sensitive to eye and neck movements (e.g., Kaufman et al., 1996) as multimodal information from vestibular, motor, and proprioception are already integrated within the vestibular nuclei (Angelaki and Cullen, 2008; Cullen and Taube, 2017). Even though these three brainstem nuclei contain a multimodal signal, lesion and recording studies confirm the importance of both NPH and SGN for the generation of HD signal (Clark et al., 2012; Butler and Taube, 2015) and navigation (Butler et al., 2017). Whether lesions to PGRNd also disrupt the HD signal remains unknown, as well as PGRNd’s role in navigational behavior.

Lesion studies have shown that disruption of the HD circuit afferent to the DTN leads to burst firing in ADN neurons (where HD cells are abundant), while disruption of DTN or areas efferent to it lead to non-burst firing in ADN or postsubiculum neurons (see Valerio and Taube, 2016 for a discussion of this point). Collectively, these studies suggest that the hypothesized attractor network that generates the HD signal is present across the connections between the DTN and LMN. While both the NPH and SGN send projections to the DTN, the SGN → DTN projections are considerably denser than the NPH → DTN projections (Graham et al., 2021; Mehlman et al., 2021), suggesting that the SGN plays a more pivotal role in generating the HD signal.

Neurons sensitive to AHV have been reported in a number of cortical areas including the retrosplenial cortex, posterior parietal cortex, medial entorhinal cortex, and the dorsal striatum (Wilber et al., 2014; Mehlman et al., 2019; Hennestad et al., 2021; Keshavarzi et al., 2022; Spalla et al., 2022). Whether the AHV signal is conveyed to each of these areas via the NPH or SGN, or alternatively via other brainstem areas, awaits further investigation. Two distinct pathways have been identified from the vestibular nuclei to the cortex – an anterior pathway through the anterior thalamic nuclei and a posterior pathway via the ventral posterior thalamus (Cullen and Taube, 2017). Interestingly, lesions of the anterior thalamus abolished the HD signal in the dorsal striatum without interfering with the AHV signal (Mehlman et al., 2019), suggesting that the striatal AHV signal reaches the striatum prior to conveying the AHV information to the anterior thalamus.

### Active vs. passive responses

A second aim of this study was to compare the properties of AHV cells during active versus passive rotations and compare their responses to previous studies in vestibular areas, which found a suppression of neuronal activity during active (volitional) rotations (Roy and Cullen, 2001, 2004; Carriot et al., 2013; Medrea and Cullen, 2013). In our relatively small sample size, we observed a variety of different responses, with some cells remaining sensitive to AHV in each brainstem nucleus, while other cells lost their AHV tuning. The diverse responses we found during passive rotation is consistent with a previous study recording AHV cells in DTN under hand-held conditions (Sharp et al., 2001b). It is also consistent with the reports cited above of variable levels of attenuation in vestibular neurons during active movement, including a small population of vestibular neurons that are minimally suppressed during active movement (Khalsa et al., 1987; Phillips et al., 1996; Shinder and Taube, 2014). While it may be argued that some AHV suppression was observed in our results during active versus passive conditions (e.g., Fig. 10 A-C), our findings indicate that these cells were still highly sensitive to AHV during active foraging.

### Relevance to computational ring-attractor models

The presence of AHV cells during active movement within NPH, SGN, and PGRNd provides support for continuous ring attractor models that use AHV cells to update directional heading (McNaughton et al., 1991; Redish et al., 1996; Zhang, 1996; Clark and Taube, 2012; Song and Wang, 2005). According to these models, each HD is conceptually placed in a ring representing every possible direction. AHV cells contribute to the attractor ring by providing information about the direction and speed of the head movement, such that when the animal turns its head, the hill of activity shifts appropriately and the peak activity of the network represents the new HD. Furthermore, some models propose a role for AHV × HD cells, where AHV cells project first to AHV × HD cells, which in turn activate HD cells to the right or to the left of the current peak activity in order to shift the activity hill in the correct direction. The finding of an AHV × HD cell within the SGN (although only one cell) lends support for models that use these types of units.

One difficulty for most continuous ring attractor models is the lack of consideration for symmetrical AHV cells. It is obvious that asymmetrical AHV cells provide information needed to push the activity hill to the right or left depending on the direction of head rotation, but it is less intuitive as to the contribution of symmetrical AHV cells. Given that symmetrical and asymmetrical cells were recorded not only within NPH, SGN, and PGRNd, but also in the DTN and LMN, it becomes important to understand how these cells contribute to the HD signal. One possibility is that symmetrical AHV cells are necessary in the calibration or fine-tuning of the HD system to prevent drift in HD hill of activity, as the initial connections following development are unlikely to be balanced perfectly (Stratton et al., 2010).

Recent work in the *Drosophila* central complex have identified circuits involved in encoding directional heading (Seelig et al., 2015; Hulse and Jayaraman, 2020). The circuit includes cells adjacent to the protocerebral bridge, referred to as P-EN neurons, that are tuned to AHV (Turner-Evans, 2017; Lyu et al., 2022). These neurons all have asymmetric tuning curves and currently no symmetric AHV cells have been identified. Given the large number of symmetric AHV cells observed in NPH and SGN, it’s not clear that the underlying neural mechanisms contributing to HD sensitivity in the *Drosophila* are ultimately going to be similar to that in a mammalian system. Certainly, a role for symmetric AHV cells must be discerned. Likewise, although few in number, the identification of offset cells in both asymmetric and inverted cells provides an interesting way to initiate a shift in an attractor style network.

### Conclusions

In summary, many cells recorded within the NPH, SGN, and PGRNd correlated with the animal’s AHV during active and, in some cases, passive rotations of the head. These findings suggest that these nuclei provide important information about head movements for the HD system, and provide further insight into why lesions to NPH (Butler and Taube, 2015) and SGN (Clark et al., 2012) disrupt HD cells within ADN.

## Acknowledgements

This work was supported by NIH grants: NS053907, NS111695. The authors wish to thank Jennifer Marcroft for her technical assistance and help with recordings, and to Roddy Grieves for comments on the manuscript. The authors have no conflict of interest to declare.

## Contributions

JRD, SSW, JEB, JST designed the experiments; JRD, SSW, JEB, CCA, KBF, AJM, NAS performed the experiments, JAG, JRD, PAL, JST analyzed the data; JAG, JRD, JST wrote the manuscript.

## Declaration of Interests

**Extended Data Figure 1**

**Included cells.** Cells that did not pass threshold criteria for correlation and slope, but were classified as AHV cells and included in the analyses. Cells 1-3 did not pass the 99^th^ percentile shuffle procedure, but cell 1 passed the 95^th^ percentile shuffle. Cells 4 and 5 did pass the 99^th^ percentile shuffle procedure. Cells 1 and 3 were in NPH, cell 2 was in PGRNd, cells 4 and 5 were in SGN. Labels for all plots, as well as CW and CCW values, are as depicted for cell 1.

**Extended Data Figure 2**

**Spike waveform widths.** 74 AHV cells + 79 non-AHV cells were recorded on a Neuralynx system that allowed analysis of spike waveforms. Histogram plots of spike width are shown above based on cell type (Left column: symmetric, asymmetric, non-AHV), brain region (middle column: NPH, SGN, PGRNd), and which hemisphere showed increased firing relative to the recorded hemisphere (right column: ipsiversive, contraversive). All plots are best characterized as gaussian in nature and there was no major differences between plots – indicating that there were no discernible bimodality in spike width distributions and thus no clear separation into putative excitatory or inhibitory populations. The dashed vertical line in each plot represents the mean spike waveform width for that group.

## References

Angelaki DE, Cullen KE (2008) Vestibular system: the many facets of a multimodal sense. Ann Rev Neurosci 31: 125–150.

Bartho P, Hirase H, Monconduit L, Zugaro M, Harris KD, Buzsaki G (2004) Characterization of neocortical principal cells and interneurons by network interactions and extracellular features. J Neurophysiol 92: 600–608.

Bassett JP, Taube JS (2001) Neural correlates for angular head velocity in the rat dorsal tegmental nucleus. J Neurosci 21: 5740–5751.

Bassett JP, Tullman ML, Taube JS (2007) Lesions of the tegmentomammillary circuit in the head direction system disrupt the head direction signal in the anterior thalamus. J Neurosci 27: 7564–7577.

Biazoli Jr CE, Goto M, Campos AM, Canteras NS (2006) The supragenual nucleus: a putative relay station for ascending vestibular signs to head direction cells. Brain Res 1094: 138–148.

Burgess N, Cacucci F, Lever C, O’Keefe J (2005) Characterizing multiple independent behavioral correlates of cell firing in freely moving animals. Hippocampus 15: 149–153.

Butler WN, Smith KS, van der Meer MAA, Taube JS (2017). The head-direction signal plays a functional role as a neural compass during navigation. Curr Biol 27: 1259–1267.

Butler WN, Taube JS (2015) The nucleus prepositus hypoglossi contributes to head direction cell stability in rats. J Neurosci 35: 2547–2558.

Cameron AC, Windmeijer FAG (1996) R-squared measures for count data regression models with applications to health-care utilization. J Business Economic Stat 14: 209–220.

Carriot J, Brooks JX, Cullen KE (2013) Multimodal integration of self-motion cues in the vestibular system: active versus passive translations. J Neurosci 33: 19555–19566.

Clark BJ, Bassett JP, Wang SS, Taube JS (2010) Impaired head direction cell representation in the anterodorsal thalamus after lesions of the retrosplenial cortex. J Neurosci 30: 5289–5302.

Clark BJ, Brown JE, Taube JS (2012) Head direction cell activity in the anterodorsal thalamus requires intact supragenual nuclei. J Neurophysiol 108: 2767–2784.

Clark BJ, Taube JS (2012) Vestibular and attractor network basis of the head direction cell signal in subcortical circuits. Front Neural Circuits 6: 7.

Clark BJ, LaChance PA, Winter SS, Mehlman ML, Butler WN, Taube JS (2023) Comparison of head direction cell firing characteristics across thalamo-parahippocampal circuitry. Hippocampus, pending revisions.

Cullen KE, Chen-Huang C, McCrea RA (1993) Firing behavior of brain stem neurons during voluntary cancellation of the horizontal vestibuloocular reflex. II. Eye movement related neurons. J Neurophysiol 70: 844–856.

Cullen KE, Taube JS (2017) Our sense of direction: progress, controversies and challenges. Nat Neurosci 20: 1465–1473.

Dale A, Cullen KE (2013) The nucleus prepositus predominantly outputs eye movement-related information during passive and active self-motion. J Neurophysiol 109: 1900–1911.

Delgado-Garcia JM, Vidal PP, Gomez C, Berthoz A (1989) A neurophysiological study of prepositus hypoglossi neurons projecting to oculomotor and preoculomotor nuclei in the alert cat. Neurosci 29: 291–307.

Escudero M, Cheron G, Godaux E (1996) Discharge properties of brain stem neurons projecting to the flocculus in the alert cat. II. Prepositus hypoglossal nucleus. J Neurophysiol 76: 1775–1785.

Fukushima K, Kaneko CRS, Fuchs AF (1992) The neuronal substrate of integration in the oculomotor system. Prog Neurobiol 39: 609–639.

Graham JA, McGuier M, Chang MS, Gundlach AC, Taube JS (2021) Projections to the lateral mammillary and dorsal tegmental nuclei arise from a small but overlapping population of neurons in the supragenual nucleus. Program No. P863.07. 2021 Abstract Viewer/Itinerary Planner. Washington DC: Society for Neuroscience. Online.

Grantyn A, Ong-Meang Jacques V, Berthoz A (1987) Reticulo-spinal neurons participating in the control of synergic eye and head movements during orienting in the cat. II. Morphological properties as revealed by intra-axonal injections of horseradish peroxidase. Exp Brain Res 66: 355–377.

Hardcastle Z, Maheswaranathan N, Ganguli S, Giocomo LM (2017) A multiplexed, heterogeneous, and adaptive code for navigation in medial entorhinal cortex. Neuron 94: 375–387.

Hayakawa T, Zyo K (1990) Fine structure of the lateral mammillary projection to the dorsal tegmental nucleus of Gudden in the rat. J Comp Neurol 298: 224–236.

Hennestad E, Witoelar A, Chambers AR, Vervaeke K (2021) Mapping vestibular and visual contributions to angular head velocity tuning in the cortex. Cell Reports 37: 110134.

Hulse BK, Jayaraman V (2020) Mechanisms underlying the neural computation of head direction. Ann Rev Neurosci 43: 31–54.

Jacob P-Y, Casali G, Spieser L, Page H, Overington D, Jeffery K (2017) An independent, landmark-dominated head-direction signal in dysgranular retrosplenial cortex. Nat Neurosci 20: 173–175.

Kaufman GD, Mustari MJ, Miselis RR, Perachio AA (1996) Transneuronal pathways to the vestibulocerebellum. J Comp Neurol 370: 501–523.

Keshavarzi S, Bracey EF, Faville RA, Campagner D, Tyson AL, Lenzi SC, Branco T, Margrie TW (2022) Multisensory coding of angular head velocity in the retrosplenial cortex. Neuron 110: 1–12.

Khalsa SB, Tomlinson RD, Schwarz DW, Landolt JP (1987) Vestibular nuclear neuron activity during active and passive head movement in the alert rhesus monkey. J Neurophysiol 57: 1484–1497.

Kitama T, Grantyn A, Berthoz A (1995) Orienting-related eye–neck neurons of the medial ponto-bulbar reticular formation do not participate in horizontal canal-dependent vestibular reflexes of alert cats. Brain Res Bull 38: 337–347.

Kubie JL (1984) A driveable bundle of microwires for collecting single-unit data from freely-moving rats. Physiol Behav 32: 115–118.

LaChance PA, Graham J, Shapiro BL, Morris AJ, Taube JS (2022) Landmark-modulated directional coding in postrhinal cortex. Science Adv 8: eabg8404.

Lannou J, Cazin L, Precht W, Le Taillanter M (1984) Responses of prepositus hypoglossi neurons to optokinetic and vestibular stimulations in the rat. Brain Res 301: 39–45.

Liu R, Chang L, Wickern G (1984) The dorsal tegmental nucleus: an axoplasmic transport study. Brain Res 310: 123–132.

Lopez-Barneo J, Darlot C, Berthoz A, Baker R (1982) Neuronal activity in prepositus nucleus correlated with eye movement in the alert cat. J Neurophysiol 47: 329–352.

Lyu C, Abbot LF, Maimon G (2022) Building an allocentric traveling direction signal via vector computation. Nature 601: 92–97.

Massot C, Schneider AD, Chacron MJ, Cullen KE (2012) The vestibular system implements a linear-nonlinear transformation in order to encode self-motion. PLoS Biol 10(7): e1001365.

McCrea RA, Gdowski GT, Boyle R, Belton T (1999) Firing behavior of vestibular neurons during active and passive head movements: vestibulo-spinal and other non-eye-movement related neurons. J Neurophysiol 82: 416–428.

McCrea RA, Horn AK (2006) Nucleus prepositus. Prog Brain Res 151: 205–230.

McFarland JL, Fuchs AF (1992) Discharge patterns in nucleus prepositus hypoglossi and adjacent medial vestibular nucleus during horizontal eye movement in behaving macaques. J Neurophysiol 68: 319–332.

McCormick DA, Connors BW, Lighthall JW, Prince DA (1985) Comparative electrophysiology of pyramidal and sparsely spiny stellate neurons of the neocortex. J Neurophysiol 54: 782–806.

McNaughton BL, Chen LL, Markus EJ (1991) “Dead reckoning,” landmark learning, and the sense of direction: a neurophysiological and computational hypothesis. J Cogn Neurosci 3: 190–202.

Medrea I, Cullen KE (2013) Multisensory integration in early vestibular processing in mice: The encoding of passive versus active motion. J Neurophysiol 110: 2704–2717.

Mehlman ML, Marcroft JL, Taube JS (2021) Anatomical projections to the dorsal tegmental nucleus and abducens nucleus arise from separate cell populations in the nucleus prepositus hypoglossi, but overlapping cell populations in the medial vestibular nucleus. J Comp Neurol 529: 2706–2726.

Mehlman ML, Winter SS, Valerio S, Taube JS (2019) Functional and anatomical relationships between the medial precentral cortex, dorsal striatum, and head direction cell circuity. I. Recording studies. J Neurophysiol 121: 350–370.

Miles FA (1974) Single unit firing patterns in the vestibular nuclei related to voluntary eye movements and passive body rotation in conscious monkeys. Brain Res 71: 215–224.

Muir GM, Brown JE, Carey JP, Hirvonen TP, Della Santina CC, Minor LB, Taube JS (2009) Disruption of the head direction cell signal after occlusion of the semicircular canals in the freely moving chinchilla. J Neurosci 29: 14521–14533.

Nishiike S, Guldin WO, Bäurle J (2000) Corticofugal connections between the cerebral cortex and the vestibular nuclei in the rat. J Comp Neurol 420: 363–372.

Paxinos G, Watson C (2005) The Rat Brain in Stereotaxic Coordinates (5th ed.). San Diego, CA: Academic.

Phillips JO, Ling I, Siebold C, Fuchs AF (1996) Behavior of primate vestibulo-ocular reflex neurons and vestibular neurons during head-free gaze shifts. Ann NY Acad Sci 781: 276–291.

Redish AD, Elga AN, Touretzky DS (1996) A coupled attractor model of the rodent head direction system. Netw Comput Neural Syst 7: 671–685.

Robinson DA (1989) Integrating with neurons. Ann Rev Neurosci 12: 33–45.

Roy JE, Cullen KE (2001) Selective processing of vestibular reafference during self-generated head motion. J Neurosci 21: 2131–2142.

Roy JE, Cullen KE (2004) Dissociating self-generated from passively applied head motion: neural mechanisms in the vestibular nuclei. J Neurosci 24: 2102–2111.

Seelig JD, Jayaraman V (2015) Neural dynamics for landmark orientation and angular path integration. Nature 521: 186–191.

Sharp PE, Blair HT, Cho J (2001a) The anatomical and computational basis of the rat head-direction cell signal. Trends Neurosci 24: 289–294.

Sharp PE, Tinkelman A, Cho J (2001b) Angular velocity and head direction signals recorded from the dorsal tegmental nucleus of gudden in the rat: implications for path integration in the head direction cell circuit. Behav Neurosci 115: 571–588.

Shinder ME, Taube JS (2011) Active and passive movement are encoded equally by head direction cells in the anterodorsal thalamus. J Neurophysiol 106: 788–800.

Shinder ME, Taube JS (2014) Resolving the active versus passive conundrum for head direction cells. Neuroscience 270: 123–138.

Song P, Wang XJ (2005) Angular path integration by moving hill of activity: A spiking neuron model without recurrent excitation of the head-direction system. J Neurosci 25: 1002–1014.

Spalla D, Treves A, Boccara CN (2022) Angular and linear speed cells in the parahippocampal circuits. Nature Comm 13: 1–13.

Stackman RW, Taube JS (1997) Firing properties of head direction cells in rat anterior thalamic neurons: dependence upon vestibular input. J Neurosci 17: 4349–4358.

Stackman RW, Taube JS (1998) Firing properties of rat lateral mammillary single units: head direction, head pitch, and angular head velocity. J Neurosci 18: 9020–9037.

Stratton P, Wyeth G, Wiles J (2010) Calibration of the head direction network: a role for symmetric angular head velocity cells. J Comput Neurosci 28: 527–538.

Taube JS (1995) Head direction cells recorded in the anterior thalamic nuclei of freely moving rats. J Neurosci 15: 70–86.

Taube JS (2007) The head direction signal: origins and sensory-motor integration. Ann Rev Neurosci 30: 181–207.

Taube JS, Muller RU, Ranck Jr JB (1990) Head-direction cells recorded from the postsubiculum in freely moving rats. II. Effects of environmental manipulations. J Neurosci 10: 436–447.

Tempia F, Ghirardi M, Dotta M, Strata P (1992) Spontaneous gaze shifts in intact head-free rats and following inferior olive and cerebellar lesions. Eur J Neurosci 4: 1239–1248.

Turner-Evans D, Wegener S, Rouault H, Franconville R, Wolff T, Seelig JD, Druckmann S, Jayaraman V (2017) Angular velocity integration in a fly heading circuit. eLife 6, e23496.

Uchino Y, Sasaki M, Sato H, Bai R, Kawamoto E (2005) Otolith and canal integration on single vestibular neurons in cats. Exp Brain Res 164: 271–285.

Valerio S, Taube JS (2016) Head direction cell activity is absent in mice without the horizontal semicircular canals. J Neurosci 36: 741–754.

Wallace DJ, Sawinski J, Greenberg DS, Sawinski J, Rulla S, Notaro G, Kerr JND (2013) Rats maintain an overhead binocular field at the expense of constant fusion. Nature 498: 65–69.

Wilber AA, Clark BJ, Forster TC, Tatsun M, McNaughton BL (2014) Interaction of egocentric and world-centered reference frames in the rat posterior parietal cortex. J Neurosci 34: 5431–5446.

Winter SS, Clark BJ, Taube JS (2015) Spatial navigation. Disruption of the head direction cell network impairs the parahippocampal grid cell signal. Science 347: 870– 874.

Yoder RM, Peck JR, Taube JS (2015) Visual landmark information gains control of the head direction signal in the lateral mammillary nuclei. J Neurosci 35: 1354–1367.

Zhang K (1996) Representation of spatial orientation by the intrinsic dynamics of the head-direction cell ensemble: a theory. J Neurosci 16: 2112–2126.

