## Extended Data Figures 1 and 2 for "Angular head velocity cells within brainstem nuclei projecting to the head direction circuit"

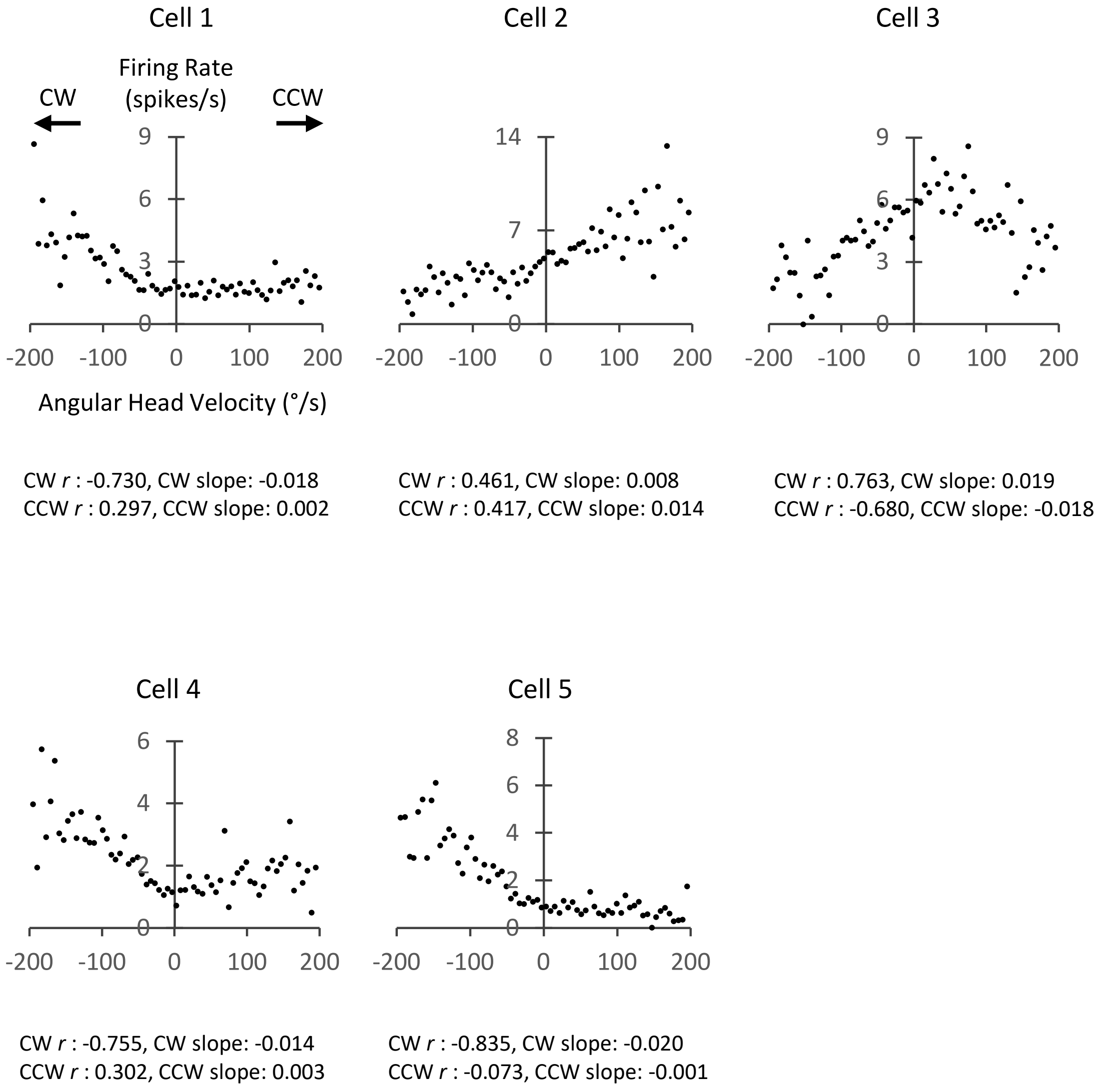


**Included cells.** Cells that did not pass threshold criteria for correlation and slope, but were classified as AHV cells and included in the analyses. Cells 1-3 did not pass the 99$^{th}$ percentile shuffle procedure, but cell 1 passed the 95$^{th}$ percentile shuffle. Cells 4 and 5 did pass the 99$^{th}$ percentile shuffle procedure. Cells 1 and 3 were in NPH, cell 2 was in PGRNd, cells 4 and 5 were in SGN. Labels for all plots, as well as CW and CCW values, are as depicted for cell 1.

### Extended Data Figure 1

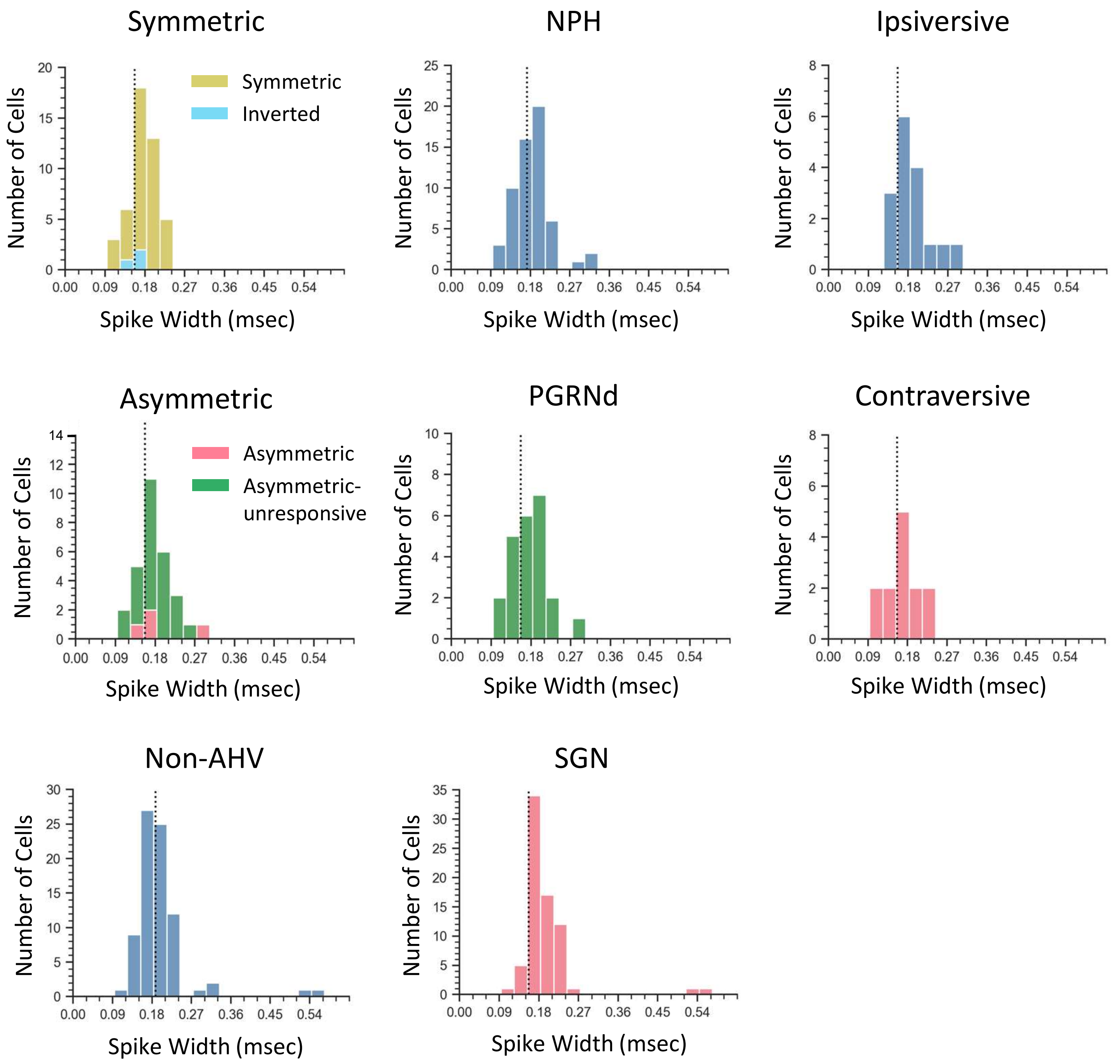


**Spike waveform widths.** 74 AHV cells + 79 non-AHV cells were recorded on a Neuralynx system that allowed analysis of spike waveforms. Histogram plots of spike width are shown above based on cell type (Left column: symmetric, asymmetric, non-AHV), brain region (middle column: NPH, SGN, PGRNd), and which hemisphere showed increased firing relative to the recorded hemisphere (right column: ipsiversive, contraversive). All plots are best characterized as gaussian in nature and there were no major differences between plots – indicating that there was no discernible bimodality in spike width distributions and thus no clear separation into putative excitatory or inhibitory populations. The dashed vertical line in each plot represents the mean spike waveform width for that group.

### Extended Data Figure 2
